# Epithelial MHC II antigen presentation dynamically informs intestinal homeostasis and injury

**DOI:** 10.64898/2026.03.18.712348

**Authors:** Vladyslav Holiar, Vladislav Rudenko, Chen Weller, Mariia Naumova, Sacha Lebon, Marco Canella, Petro Busko, Avital Sarusi-Portuguez, Tali Shalit, Aviya Habshush Menachem, Idan Adir, Zachary Petrover, Polina Greenberg, Corine Katina, Polina Gradchenko, Barak Toval, Nissan Yissachar, Irit Sagi, Eldad Tzahor, Yishai Levin, Yardena Samuels, Moshe Biton

**Affiliations:** Department of Immunology and Regenerative Biology, Weizmann Institute of Science, Rehovot, 7610001, Israel; Department of Molecular Cell Biology, Weizmann Institute of Science, Rehovot 7610001, Israel; Department of Life Science Core Facilities, Weizmann Institute of Science, Rehovot 7610001, Israel; de Botton Institute for Protein Profiling, the Nancy and Stephen Grand Israel National Center for Personalized Medicine, Weizmann Institute of Science, Rehovot, 7610001, Israel; The Goodman Faculty of Life Sciences, Bar-Ilan University, Ramat-Gan, Israel; Bar-Ilan Institute of Nanotechnology and Advanced Materials, Bar-Ilan University, Ramat-Gan, Israel

**Keywords:** 1. Intestinal epithelial cells, 2. MHC class II ligandome, 3. Immunopeptidomics, 4. CD4□ T cell responses, 5. Peripheral tolerance, 6. Damage-antigens

## Abstract

The intestinal epithelium plays a pivotal role in balancing immune tolerance and inflammation, yet how it communicates tissue state to the adaptive immune system remains unclear. Here, we show that intestinal epithelial cells (IECs) encode tissue identity and injury into the major histocompatibility complex class II (MHC II) ligandome. We employed integrated single cell transcriptomics, quantitative proteomics, and high-depth *in vivo* immunopeptidomics to map the MHC class II self-peptidome of the mouse small intestine across epithelial and immune compartments. Mature enterocytes and intestinal stem cells (ISCs) emerged as the dominant epithelial antigen-presenting cells (APCs), displaying a compartmentalized repertoire of endogenous self-immunopeptides reflecting epithelial differentiation and function. Disruption of epithelial MHC II expression led to loss of antigenic compartmentalization, immune infiltration, extracellular matrix remodeling, and emergence of inflammation-associated immune ligands, demonstrating that epithelial MHC II is required to maintain homeostasis. Functionally, a subset of ISC-derived self-immunopeptides preferentially promotes regulatory CD4□ T cell responses, linking epithelial antigen presentation and peripheral tolerance. During gut inflammation, the epithelial MHC II landscape shifted toward damage-associated antigens. Together, these findings establish epithelial MHC II presentation as a context-dependent tissue-immune communication system that promotes tolerance in homeostasis and alerts to tissue injury during inflammation.

## Introduction

The gut is a highly specialized tissue that supports nutrient and vitamin absorption while simultaneously balancing immune activation and tolerance. Central to this regulation is the adaptive immune system, which is guided by the presentation of self and non-self antigens by antigen-presenting cells (APCs) through major histocompatibility complex (MHC) molecules ^1–3^. MHC II machinery is primarily expressed by professional APCs, such as B cells, dendritic cells (DCs), and macrophages. The ability of professional APCs to induce and maintain peripheral tolerance to self-proteins and foreign antigens in the gut has long been recognized and remains a major research interest in the field ^4–9^. However, less is known about the role of MHC II in intestinal epithelial cells (IECs).

Previous studies have recognized IECs as non-conventional APCs, capable of expressing the MHC II machinery ^10–13^. Nevertheless, there is a long-standing debate about the role of MHC II on IECs. While some studies suggest a tolerogenic role of epithelial MHC II under steady-state conditions or in the context of gut inflammation ^14–16^, others indicate a pro-inflammatory role in gut pathologies, including graft-versus-host disease (GvHD) and IBD ^17,18^. Our previous study demonstrated that intestinal stem cells (ISCs) interact with CD4^+^ T cells *in vivo*. This study highlighted the importance of epithelial-T cell interactions for proper ISC function and suggests a role for ISCs in shaping the CD4^+^ T landscape ^14^. In addition, it has been shown that epithelial cells expressing the hemagglutinin (HA) induce a regulatory phenotype in HA-specific T cells, and that antigen-driven interaction between IECs and CD4^+^ T cells expands Foxp3^+^ T regulatory (Treg) cells, independent of local DCs ^19,20^. However, a direct demonstration of IEC presentation of endogenous epithelial self-antigens that enforces tolerance has not yet been demonstrated and remains largely inferred.

In the gut, there is a strong association between genetic variability in MHC II loci and susceptibility to celiac or inflammatory bowel diseases (IBD) ^21,22^. Despite this, the repertoire and nature of peripheral self-antigen presentation by gut APCs, either immune or epithelial cells, remain poorly defined. Recent advances in mass spectrometry (MS) have enabled the identification of MHC I and II immunopeptides, uncovering self or neoantigens associated with cancer and, more recently, with certain autoimmune diseases ^23–27^. Immunopeptidomics studies in autoimmune diseases have shown that the MHC II repertoire is primarily composed of self-proteins, with MHC polymorphisms influencing the diversity and stability of the presented epitopes ^23,28^. In pancreatic islets, MHC II presentation is strongly enriched for proinsulin-derived self-peptides, providing a mechanistic link between tissue-derived antigens and autoreactive T cell responses in type 1 diabetes ^29^. These findings suggest that tissue-resident APCs, by virtue of their local protein environment, display abundant self-antigens that support homeostatic immune tolerance while triggering an immune response during inflammation.

Here, we establish a critical role for epithelial MHC class II in maintaining intestinal homeostasis. We identify mature enterocytes and ISCs as epithelial APCs. To define their antigens, we comprehensively map the epithelial MHC II self-peptidome and identify a large repertoire of epithelial-specific self-peptides that are distinct from those presented by lamina propria immune cells. By integrating transcriptomic, proteomic, and immunopeptidomics analyses, we show that MHC II ligands are highly compartmentalized, with villus- and crypt-specific peptides reflecting the underlying cellular identity and functional state of the epithelium. Notably, a subset of crypt-derived peptides originating from ISC-associated proteins preferentially promotes tolerogenic CD4□ T cell responses, indicating a role for ISC-specific self-antigens in peripheral tolerance. Finally, using a DSS-induced colitis model, we demonstrate that gut inflammation reshapes the epithelial MHC II ligandome, leading to the emergence of inflammation-associated immunopeptides alerting to tissue damage. Together, these findings reveal that epithelial MHC II presentation encodes tissue state into a compartment-specific self-antigen landscape.

## Results

### Identifying MHC II-expressing cells in the gut

Major histocompatibility complex class II (MHC II) expression by intestinal epithelial cells (IECs) has been implicated in both immune tolerance and T cell activation, yet the mechanisms underlying these opposing outcomes remain unresolved ^10,14,15,17,18,30^. While epithelial and immune compartments of the gut are known to express MHC II, it is unclear which IEC subsets contribute to antigen presentation and whether epithelial MHC II is broadly distributed or restricted to specific epithelial states. To address this, we performed flow cytometric analysis using the Lgr5-CreER-eGFP reporter mouse, which enabled the identification of intestinal stem cells (ISCs) ^31^. We isolated IECs from villi, which contain differentiated epithelial cells, and from crypts of Lieberkühn, which harbor progenitors and intestinal stem cells (ISCs), alongside lamina propria (LP) immune cells (**Figures 1A-1C** and **S1A**). We detected robust expression of the MHC II subunit H2-Ab1 across all compartments, with LP antigen-presenting cells (APCs) exhibiting approximately fivefold higher surface abundance than ISCs (**Figure 1C**). To resolve epithelial MHC II expression at cellular resolution, we sorted MHC II□ and MHC II□ IECs using fluorescence-activated cell sorting (FACS), followed by bulk RNA sequencing (**Figure S1B**). Deconvolution analysis against published single-cell RNA-sequencing (scRNA-seq) datasets ^14,32^ revealed that villus IECs were dominated by enterocytes, which were similarly represented in MHC II□ and MHC II□ fractions, whereas crypt IECs displayed greater cellular diversity, with ISCs and transit-amplifying (TA) cells strongly enriched in the MHC II population (**Figures 1D**, **1E**, **S1C-S1D**, and **Table S1**). In contrast, secretory lineages, including goblet, enteroendocrine, and tuft cells, were preferentially excluded from the MHC II□ epithelial fraction in both villus and crypt compartments (**Figures 1E**, **S1G,** and **S1H**). The same analysis was done on MHC II□ and MHC II□ immune cells to identify MHC II^+^ immune cells. MHC II samples were dominated by B cells and macrophages, whereas CD4^+^ T cells comprised a considerable portion of MHC II□ samples (**Figures S1E** and **S1F**). In addition, gene ontology (GO) analysis identified 713 terms between the MHC II□ and MHC II□ populations (q <0.05) in the villus compartment, with 456 of them characterizing the MHC II□ villus population (**Figure S1G**). Key GO processes suggested for enrichment of mature enterocytes gene expression associated with lipid metabolism in MHC II^+^ compared to MHC II^-^ cells, such as apolipoprotein A (*Apoa*) family and Fatty-Acid-Binding Protein 2 (*Fabp2*) (**Figures S1G** and **S1H)**. On the other hand, MHC II□ epithelial cells from villus and crypt displayed GO terms related to peptide and hormone secretion, such as gastric inhibitory polypeptide (*Gip*) and Cholecystokinin (*Cck*), characterizing the secretory lineage of the small intestine (SI) (**Figures S1G** and **S1H**). Together, these data reveal that MHC II expression is not a generic feature of the intestinal epithelium but is instead restricted to absorptive and stem-cell compartments, providing a cellular framework for reconciling the divergent immunological roles attributed to epithelial MHC II.

**Figure 1.**
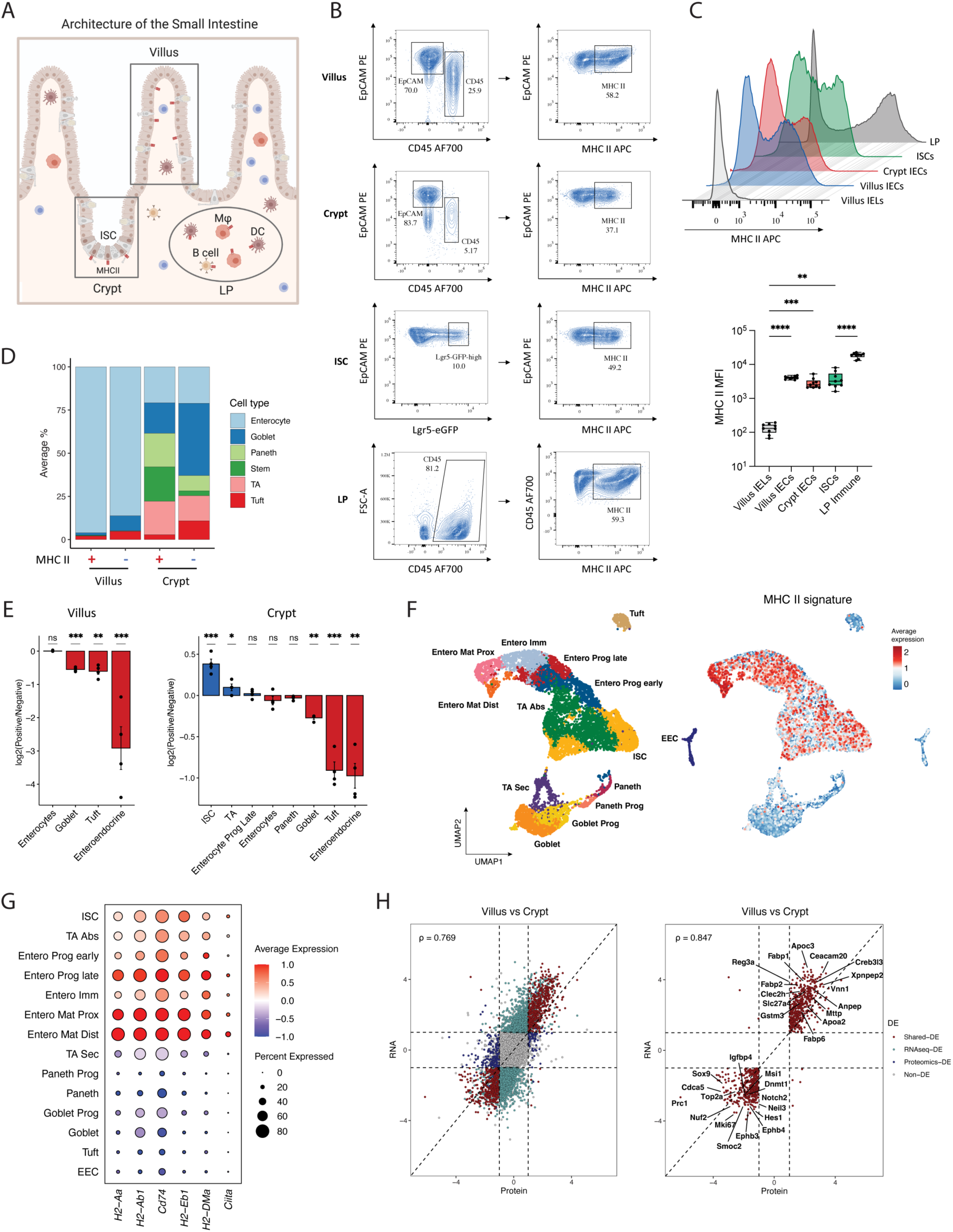
Identifying MHC II-expressing cells in the gut. **(A)** Schematic representation of the small intestine (SI) architecture. **(B)** Representative flow cytometry analysis of MHC II expression in IECs (EpCAM^+^ CD45^-^) villus and crypt, Lgr5^+^ ISCs, and CD45^+^ LP immune cells, with percentages on each panel indicating the positive fraction within the current gate. **(C)** Median fluorescence intensity of MHC II expression in villus intraepithelial lymphocytes (IELs) (white), villus (blue) and crypt (orange) IECs, ISCs (green) and LP immune cells (grey). *n* = 9 mice; statistic by repeated-measures one-way ANOVA with Geisser-Greenhouse correction, followed by Šídák-adjusted two-sided post-hoc tests. \*\**P* < 0.01 \*\*\**P* < 0.001, \*\*\*\**P* < 0.0001. **(D, E)** Bulk RNA-seq of MHC II⁺ and MHC II⁻ EpCAM^+^ CD45^-^ cells from villus and crypt of WT mice (*n* = 4). **D**, Averaged epithelial cell-type proportions inferred by the CDSeqR R package ^57^ (**Methods**) and annotated using known markers ^32^. **E**, Ratio of MHC II⁺ to MHC II⁻ epithelial signature scores per sample (**Methods**) plotted as log₂(MHC II⁺/MHC II^-^). Cell signatures curated from ^32^. Statistical significance determined by two-sample *t-tests* with Benjamini-Hochberg adjustment, \**P* < 0.05, \*\**P* < 0.01, \*\*\**P* < 0.001. **(F)** UMAP presentation of scRNA-seq of SI epithelial cells (*n* = 10,917 cells) colored by cell type (left panel) or by mean expression of the MHC II signature (right panel; **Methods**). EEC, enteroendocrine; Entero, enterocyte; Prog, progenitor; Mat, mature; Abs, absorptive; Sec, Secretory; Imm, Immature; TA, transit amplifying. **(G)** Higher expression of the MHC II gene signature in mature enterocytes, Tas, and ISCs. Dot size represents the percentage of cells expressing each gene in each cluster, and dot color indicates the mean z-scored expression across clusters. Cell subsets are on the Y-axis, genes on the X-axis. **(H)** High correlation between RNA and protein expression levels across epithelial compartments. Scatter plot of log₂ fold change at the protein (x-axis) versus RNA (y-axis) levels in MHC II⁺ epithelial cells between villus and crypt; dashed lines mark log₂FC = 1. Spearman’s ρ is reported for all differentially expressed (DE) genes/proteins (left) and only shared DE genes/proteins (right).

To define MHC class II-expressing epithelial subsets at single cell resolution, we performed single-cell RNA sequencing of the small intestinal epithelium using the 10x Genomics platform (**Figure 1F**). After quality control, we retained 10,917 epithelial cells from an initial 16,451 profiles, with a median of 2,908 genes and 8,385 unique transcripts detected per cell (**Methods**). Unsupervised graph-based clustering identified 14 transcriptionally distinct epithelial populations, which were annotated based on differentially expressed genes (DEG) and established lineage markers (**Figures 1F**, **S2A**, and **Table S2**). Across this landscape, MHC II expression was broadly confined to the absorptive lineage, spanning ISCs, TAs, enterocyte progenitors, and immature and mature enterocytes (**Figures 1F** and **1G**). Notably, mature proximal and distal enterocytes exhibited substantially higher MHC II expression than their immature counterparts, consistent with bulk RNA-sequencing data showing enrichment of lipid metabolic programs in MHC II□ villus epithelial cells (**Figures S1G** and **S2B**). In contrast, secretory lineages, including enteroendocrine, goblet, and tuft cells and their progenitors, displayed minimal to undetectable MHC II signature expression (**Figures 1F** and **1G**), indicating that MHC II expression is selectively associated with absorptive and stem-cell epithelial states.

To validate MHC class II-expressing epithelial identities at the protein level, we performed quantitative LC-MS/MS proteomic profiling of MHC II villus and crypt IECs alongside MHC II□ LP immune cells (**Methods** and **Figures S2C-S2D**). Across the three compartments, we identified the expression of MHC II-related proteins (**Figure S2E**). Overall, we detected 7,622 unique proteins, with protein abundance patterns showing strong concordance with bulk RNA-seq data from MHC II□ populations (**Figure S2D-S2F** and **Table S3**). Differential protein expression in villus and crypt MHC II epithelial cells closely mirrored transcriptional differences, with a high correlation between shared DE genes and proteins (ρ = 0.847; **Figures 1H** and **S2D**). Proteomic analysis of villus MHC II□ IECs revealed enrichment of mature proximal and distal enterocyte markers, including Fabp1, Reg3a, Fabp6, and Clec2h, consistent with their differentiated absorptive identity. In contrast, crypt MHC II□ IECs were enriched for ISC-associated proteins (Smoc2, Ephb3, Igfbp4) and proliferation markers (Mki67, Nuf2, Prc1), confirming stem and progenitor transcriptional profile (**Figures 1H** and **S2D**). Comparison of MHC II□ epithelial and immune compartments revealed a lower, yet significant, concordance in shared DE genes and proteins (ρ = 0.607), with epithelial samples dominated by enterocyte and ISC markers and immune samples enriched for canonical APC markers, including Lsp1, Itgax, and Jchain (**Figures S2D** and **S2G**). Together, these integrated transcriptomic and proteomic analyses establish mature enterocytes and ISCs as the major epithelial sources of MHC II expression in the small intestine and demonstrate that epithelial MHC II identity is robustly encoded at both the RNA and protein levels.

### Characterizing the MHC II-associated self-ligandome of the gut

The gut is a central site of immune education and peripheral tolerance ^3,33,34^, yet the composition of the MHC II ligandome in intestinal tissues remains poorly defined, in particular, the identity of the epithelial-derived class II peptides. To comprehensively identify the diversity and landscape of small intestinal antigens, we developed an *in vivo* MHC II immunopeptidomics assay of the gut (**Figure 2A**). Briefly, we modified the epithelial mechanical extraction protocol followed by immune isolation used for single cell RNA-seq analysis to purify the villus, crypt, and immune compartments of the small intestine followed by MHC II immunoprecipitation and high-sensitivity LC-MS/MS acquisition ^35^ (**Methods**). The resulting purification exhibited high compartmental purity, with minimal epithelial contamination in the immune fraction and strong enrichment of epithelial cells in villus and crypt preparations (**Figures S3A-S3D**). Across all replicates, we identified more than 10,000 peptides at 1% false discovery rate (FDR), retaining 9,207 unique intestinal sequences after quality control (**Figure 2B**). Restricting analysis to peptides predicted to bind the I-A□ allele yielded a high-confidence set of 5,180 MHC II immunopeptides (Rank≤10, NetMHCIIpan4.3, **Methods**) ^36^ (**Figures 2B**, **S4A-S4C**, and **Table S4**). As expected, lamina propria immune cells presented the largest number of ligands (3,737 peptides), whereas epithelial compartments contributed a comparable number of antigens (2,634 peptides), with the crypt alone accounting for 2,143 peptides (**Figures 2C** and **S4A**). Notably, despite high MHC II expression, villus epithelial cells presented fewer peptides (887 total) and exhibited a lower fraction of predicted binders, whereas crypt and immune compartments were enriched for high-affinity ligands. Peptide length distributions, binding ranks, and consensus motifs were highly consistent across compartments, matching canonical I-A□ binding characteristics ^37,38^ (**Figures S4B-S4E**), confirming the robustness of the dataset. Together, these analyses establish a comprehensive atlas of the intestinal MHC II ligandome and reveal that small intestinal epithelial cells, particularly within the crypt compartment, actively process and present a diverse repertoire of endogenous class II self-peptides *in vivo*.

**Figure 2.**
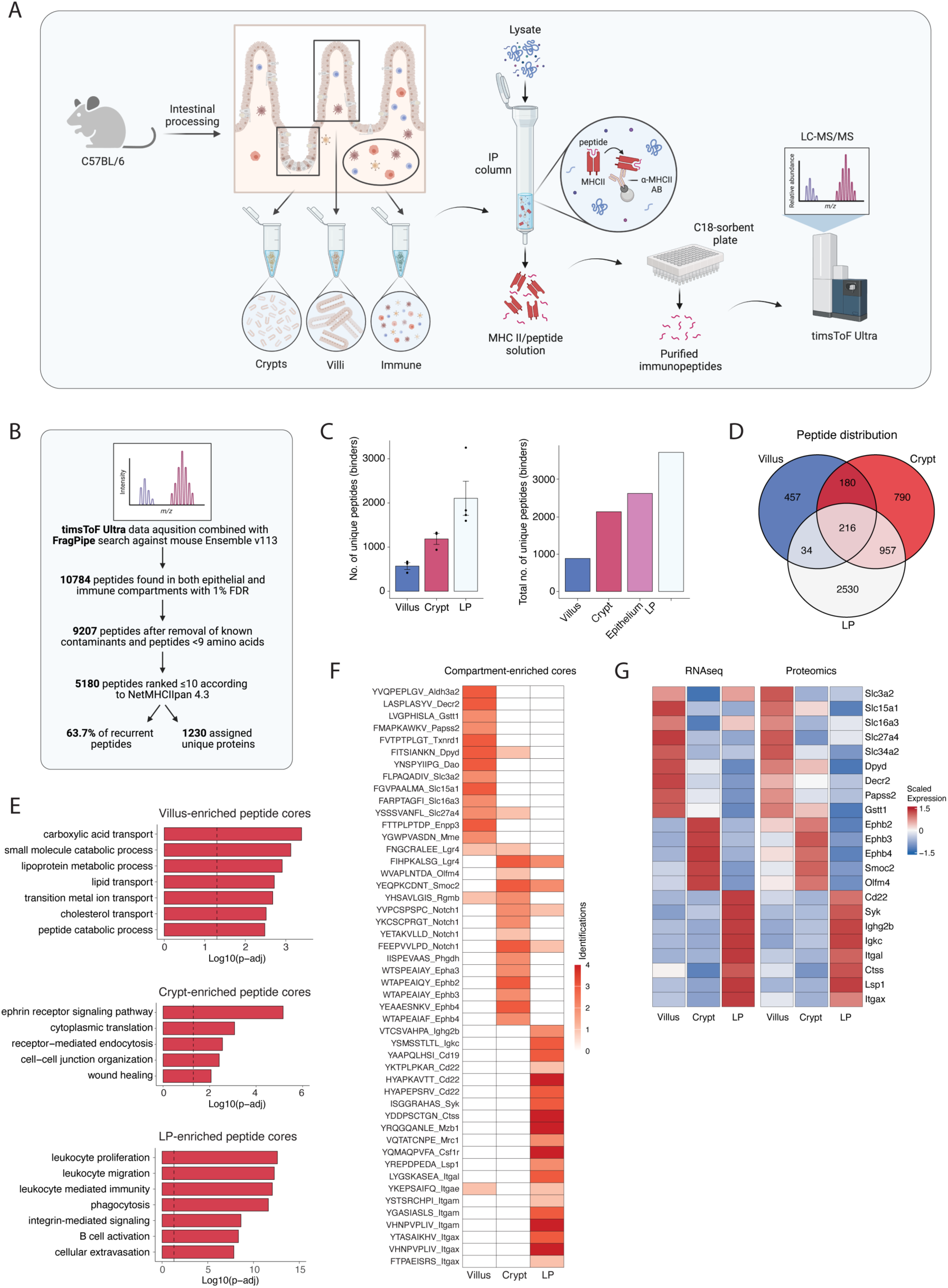
MHC II-peptidomics reveals compartmentalization of the intestinal ligandome. **(A)** Schematic of the MHC II-peptidomics experimental workflow. The small intestine (SI) of C57BL/6J mice was harvested, and villi, crypts, and LP immune cells were isolated and subjected to MHC II-bound peptide purification, followed by LC-MS/MS. **(B)** Schematic of the pipeline deployed to identify MHC II-bound peptides. Numbers indicate data pooled across villus, crypt, and LP compartments. **(C)** Number of unique MHC II-bound peptides per compartment after quality control. The left panel shows mean ± SEM with individual samples overlaid; the right panel shows the total number of unique sequences per compartment, with Epithelium representing villus and crypt compartments combined. **(D)** Distribution of MHC II-bound peptide repertoires of the SI across the villus, crypt, and LP compartments. Numbers in each region indicate the number of unique peptide sequences after filtering. Biological replicates were pooled. **(E)** Gene Ontology (GO) enrichment analysis of source proteins mapped from compartment-enriched peptide cores (from top to bottom: villus, crypt, and LP) (**Methods**). Enriched peptide cores were mapped to source proteins and analyzed per compartment using over-representation analysis (clusterProfiler; Benjamini-Hochberg FDR). Bars show log₁₀(q-value) and dashed lines mark *q* = 0.05. Representative significant terms are shown. **(F)** Compartment-enriched peptide cores followed by their source protein faithfully recapitulate *in vivo* biological functions; columns are grouped by SI compartment. Each tile indicates the number of biological replicates in which the peptide core was detected in the given compartment, following the same enrichment rule as in (**E)**. **(G)** RNA (left) and protein (right) expression levels matched to compartment-enriched peptide cores. Columns are grouped by SI compartment and assay (RNA, bulk RNA-seq; protein, proteomics). Tiles show row-wise z-scores of the log₂-normalized group means, calculated separately for RNA and protein. Genes/proteins were selected by mapping enriched peptide-carrying proteins to gene symbols.

The intestinal MHC class II immunopeptidome displayed substantial diversity, with many source proteins represented by a single peptide and a dominant contribution from extracellular, plasma membrane, and vesicular proteins, which are canonical reservoirs of class II ligands ^39,40^ (**Figures S4G** and **S4H**). Consistent with the distinct biological functions of villus, crypt, and immune compartments, MHC II-bound peptides showed strong compartment specificity at the levels of peptide sequences, binding cores, and source proteins (**Figures 2D** and **S4F**). Applying stringent criteria to define compartment-enriched ligands (**Methods**), we identified 2,530 immune-specific peptides derived from 478 proteins, 790 crypt-specific peptides from 129 proteins, and 457 villus-specific peptides from 67 proteins (**Figures 2D**, **S4F**, and **Table S4**). GO analysis revealed that compartment-enriched peptides faithfully reflected the core biological programs of their compartment of origin (**Figures 2E** and **2F**). Villus epithelial MHC II ligands were predominantly derived from proteins involved in digestion and nutrient absorption, with a strong enrichment for lipid metabolic pathways, consistent with MHC II expression in mature enterocytes (**Figures 1F**, **1G**, **2E**, and **2F**). In contrast, crypt epithelial peptides were enriched for ISC-enriched proteins, including Rgmb, Smoc2, and Olfm4 ^32^, and showed prominent representation of ephrin receptor signaling pathway critical for stem cell maintenance ^41^ (**Figures 2E**, **2F**, **S5A**, and **S5B**). The immune MHC II ligandome was dominated by peptides derived from macrophage, dendritic cell, and B cell proteins, including Mrc1, Csf1r, Itgax, Cd19, and Syk (**Figures 2E** and **2F**). Across compartments, class II peptides closely mirrored transcriptomic and proteomic expression patterns, indicating that antigen processing broadly samples the steady-state proteome of APCs and imprints this information onto the MHC II ligandome (**Figures 2F** and **2G**). Together, these findings establish the MHC II immunopeptidome as a cellular barcode that encodes both APC identity and its anatomical localization and biological function *in vivo*.

### Validation of epithelial-specific class II self-peptide landscape

To validate the presence of epithelial self-peptides and test whether epithelial MHC class II actively licenses tissue-specific antigen presentation, we employed an inducible epithelial-specific MHC II knockout model (Villin-CreERT2 crossed to H2-Ab1^fl/fl^ - MHC II^ΔIEC^, **Methods**; **Figure 3A**). Tamoxifen administration resulted in epithelial MHC II ablation without detectable changes in the frequency of LP MHC II macrophages or dendritic cells (**Figures S5A** and **S5B**). MHC II immunopeptidomics profiling of villus, crypt, and LP compartments from MHC II^ΔIEC^ mice revealed an approximately tenfold loss of villus- and crypt-specific class II ligands, demonstrating that epithelial MHC II is required to license the presentation of epithelial self-antigens *in vivo* (**Figures 3B** and **3C**). Despite this profound loss of epithelial-derived ligands, the total number of peptides recovered from epithelial fractions was reduced only by two-fold, most prominently in the crypt, indicating that immune cell presentation replaced epithelial antigenic space when epithelial MHC II is absent (**Figures 3B**, **S5C-S5E**, and **S5I**).

**Figure 3.**
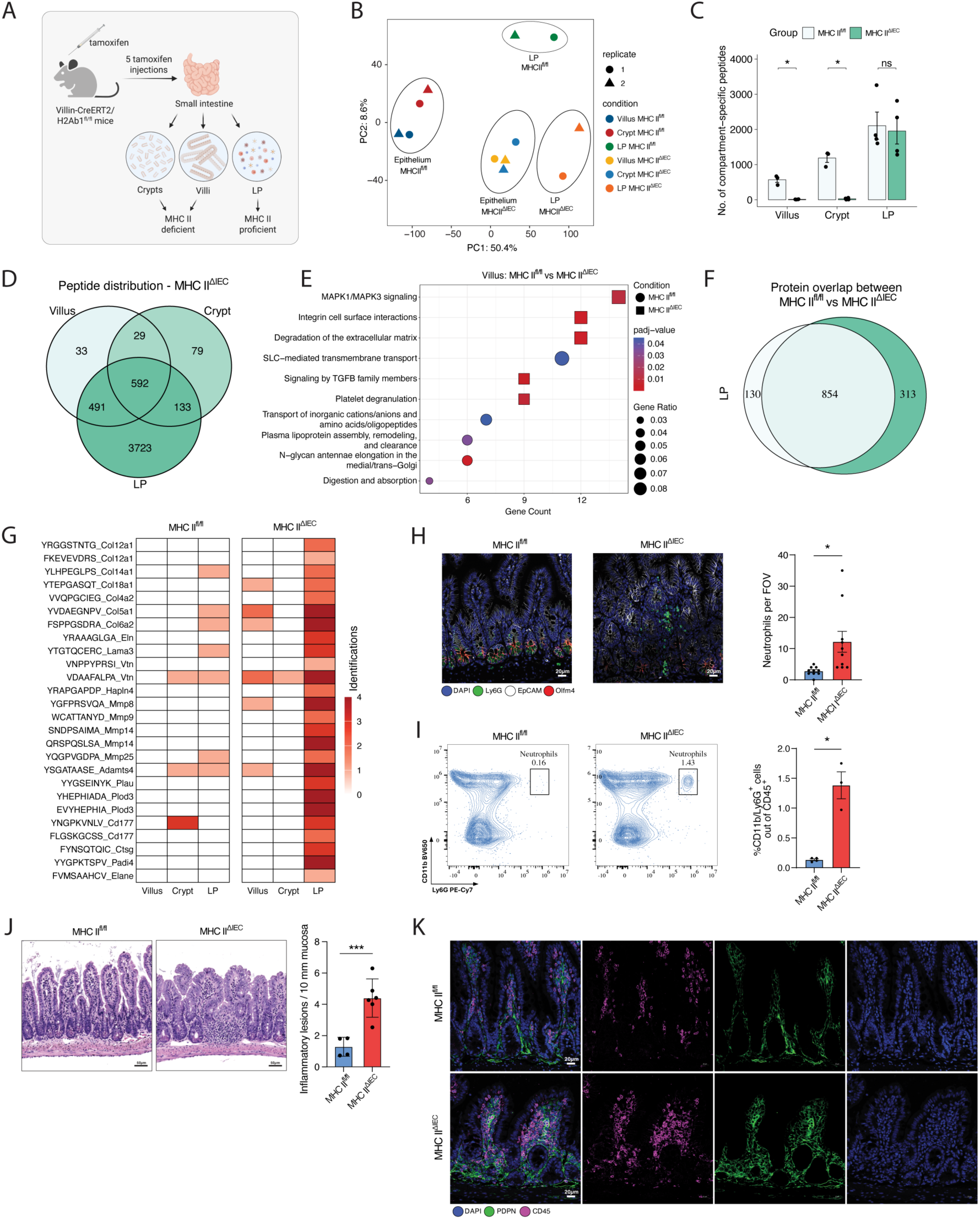
Confirming the origin of IEC-presented class II peptides. **(A)** Schematic of the experimental workflow. MHC II deletion on IECs (MHC II^ΔIEC^) was induced by tamoxifen administration. Villus, crypt, and LP compartments of the small intestine (SI) were isolated and subjected to MHC II-bound peptide purification followed by LC-MS/MS. Data represent *n = 4* mice unless specified otherwise. **(B)** Loss of immunopeptidome compartmentalization shown by Principal Component Analysis (PCA) of immunopeptides from the villus, crypt, and LP immune compartments in MHC II^ΔIEC^ compared to MHC II ^fl/fl^ mice. Axes indicate variance explained; *n* = 2 mice per genotype. **(C)** Number of compartment-specific MHC II-bound peptides in MHC II^fl/fl^ and MHC II^ΔIEC^ mice. Data are shown as mean ± SEM, and statistical analysis was performed using multiple unpaired *t-tests* with Holm-Šídák correction (α = 0.05). \**P* < 0.05. **(D)** MHC II ligandome shift represented by a Venn diagram showing the distribution of the discovered peptides across intestinal compartments of MHC II^ΔIEC^. Counts are pooled across replicates after filtering. **(E)** Cellular reprogramming identified using Reactome pathway enrichment analysis of unique source proteins from the villus peptidome of MHC II^fl/fl^ compared to MHC II^ΔIEC^ mice by the ReactomePA v1.16.2 R package ^58^. Significance was assessed using Benjamini-Hochberg correction, *q* < 0.05. **(F)** The overlap between the source proteins of the LP immunopeptidome in MHC II^fl/fl^ and MHC II^ΔIEC^ mice is shown in an area-proportional Venn diagram. Counts are pooled across replicates. **(G)** Identified peptide cores from ECM-remodeling proteins. Rows are cores followed by source proteins. Identifications represent the number of biological replicates in which a peptide core was detected (per compartment). **(H)** Immunofluorescence analysis of distal SI sections from MHC II^fl/fl^ and MHC II^ΔIEC^ mice. Representative images (left panels) show Ly6G (green), E-Cadherin (white), Olfm4 (red), and nuclei (blue). Scale bar 20 µm. Quantification of Ly6G^+^ cells per field of view (FOV; 20x), where each point represents individual FOV (Right panel). Statistical analysis was performed using a two-tailed unpaired Student’s *t-test* (*n* = 2 mice per group); data shown as mean ± SEM. \**P* < 0.05. **(I)** Neutrophil infiltration in MHC II^ΔIEC^ mice compared to MHC II^fl/fl^ controls assessed by flow cytometry analysis of the distal SI (left panels). Percentage of CD11b^+^ Ly6G^+^ neutrophils out of CD45^+^ cells displayed as mean ± SEM (right panel). Statistical analysis was done using a two-tailed unpaired *t-test* with Welch’s correction on *n* = 4 mice of MHC II^fl/fl^ vs. *n* = 3 mice of MHC II^ΔIEC^. \**P* < 0.05. **(J)** H&E staining of distal SI sections from MHC II^fl/fl^ and MHC II^ΔIEC^ mice (left panels) and quantification of inflammatory lesions per 10 mm of mucosal layer shown as mean ± SEM (right panel). Scale bar: 50µm. Statistical significance was assessed using two-tailed unpaired *t-test*. **(K)** Fibroblast accumulation in MHC II^ΔIEC^ mice was identified by immunofluorescence staining of distal SI sections stained for PDPN (green) and CD45 (purple). Nuclei stained with DAPI (blue). Scale bar: 20µm.

The collapse of epithelial-specific immunopeptides was evident at the levels of peptide sequences, binding cores, and source proteins, and was captured by principal component analysis (PCA), which revealed a loss of segregation between epithelial and immune immunopeptidomes upon epithelial MHC II deletion (**Figures 3B-D**, **S5F-S5I**, **S6A**, **S6B**, and **Table S4**). Consistent with this loss of epithelial antigenic licensing, analysis of peptide source proteins revealed a pronounced shift of the immunopeptidome. Whereas villus-derived ligands in control mice reflected absorptive epithelial programs, villus fractions from MHC II^ΔIEC^ mice were dominated by immune-associated antigens with strong enrichment for extracellular matrix (ECM) remodeling pathways and canonical immune-APC signatures (**Figures 3E** and **S5I**). A similar shift was observed in the crypt compartment, where epithelial stem cell programs were lost, and immune interaction pathways became predominant (**Figures S5I** and **S6A**). At the level of precursor proteins, the immune immunopeptidome of MHC II^ΔIEC^ mice largely overlapped with controls but exhibited increased diversity, with peptides derived from an additional 313 source proteins (**Figures 3F**, **3G**, **S5G**, and **S5H**). These newly presented peptides were enriched in proteins involved in ECM organization, maintenance, and remodeling, including structural components (collagens, elastin [Eln], laminin α3 [Lama3]) and ECM-modifying enzymes (Mmp14, Plau, Plod3), characteristic of activated fibroblasts ^42^ (**Figures 3G** and **S6B**). In parallel, we detected peptides originating from neutrophil-associated proteins, including those derived from neutrophil-expressed matrix metalloproteinases (Mmp8, Mmp9), the transmigration marker Cd177, and neutrophil elastase (Elane) ^43–45^, indicative of inflammatory tissue remodeling (**Figure 3G**). Guided by these antigenic signatures, we directly assessed tissue composition and confirmed increased fibroblast accumulation, pronounced neutrophil infiltration, and a higher incidence of inflammatory lesions in the small intestine of MHC II^ΔIEC^ mice by IFA and flow cytometry (**Figures 3H-3K** and **S6C-S6E**). Together, we validated the identification of epithelial-specific class II self-antigens using genetic ablation of MHC II machinery in IECs. These findings indicate that epithelial MHC II preserves compartmentalized epithelial antigenic identity and may suggest that epithelial class II self-antigens contribute to maintaining immune homeostasis.

### Determining the immunomodulating potential of the stem cell-specific class II self-antigens

Having established that IECs present endogenous self-antigens on MHC class II in a manner that reflects cellular identity, we next asked whether these epithelial-derived peptides exert immunomodulatory functions. We previously showed that ISCs can engage CD4□ T cells *in vivo*, including Tregs ^14^. To directly test the functional impact of epithelial self-antigen presentation, we focused on ISC-derived class II peptides identified in our immunopeptidomics analysis and assessed their capacity to shape CD4□ T cell responses using an immunogenicity assay (**Figure 4A** and **Methods**). Because antigen-specific T cells recognizing self-peptides are rare, mice were immunized twice with a pool of 11 ISC-derived class II self-peptides (20 μg/peptide) selected from well-established ISC proteins, including ephrin receptors, Notch1, Rgmb, Olfm4, and Phgdh ^32,46^, emulsified in complete Freund’s adjuvant (CFA) ^47^, with adjuvant-only immunized mice serving as controls (**Figures 4A** and **4B**). Twenty days after immunization, splenocytes were restimulated *ex vivo* with the peptide pool, and cytokine secretion was quantified. Peptide-immunized mice exhibited robust antigen-dependent secretion of interleukin-2 (IL-2) and interferon-γ (IFNγ), together with a significant increase in IL-10 production, whereas no such responses were detected in control cultures (**Figure 4C**). Notably, a modest but reproducible elevation in IL-10 secretion was observed even in the absence of *ex vivo* restimulation with peptide-mix, suggesting a durable tolerogenic imprint despite the use of a strong inflammatory adjuvant. To define the CD4□ T cell subsets responding to epithelial self-antigens, we analyzed lineage-defining transcription factors by flow cytometry. Peptide immunization induced a modest increase in T-bet□ Th1 cells but was dominated by a pronounced expansion of FOXP3□ Tregs, accompanied by elevated expression of T cell anergy markers (**Figures 4D**, **4E**, **S7A**, and **S7B**). These data indicate that ISC-derived class II self-peptides can elicit antigen-specific CD4□ T cell responses that are dominated by Treg expansion and anergy-associated programs (**Figures 4C**, **4E**, **S7A**, and **S7B**).

**Figure 4.**
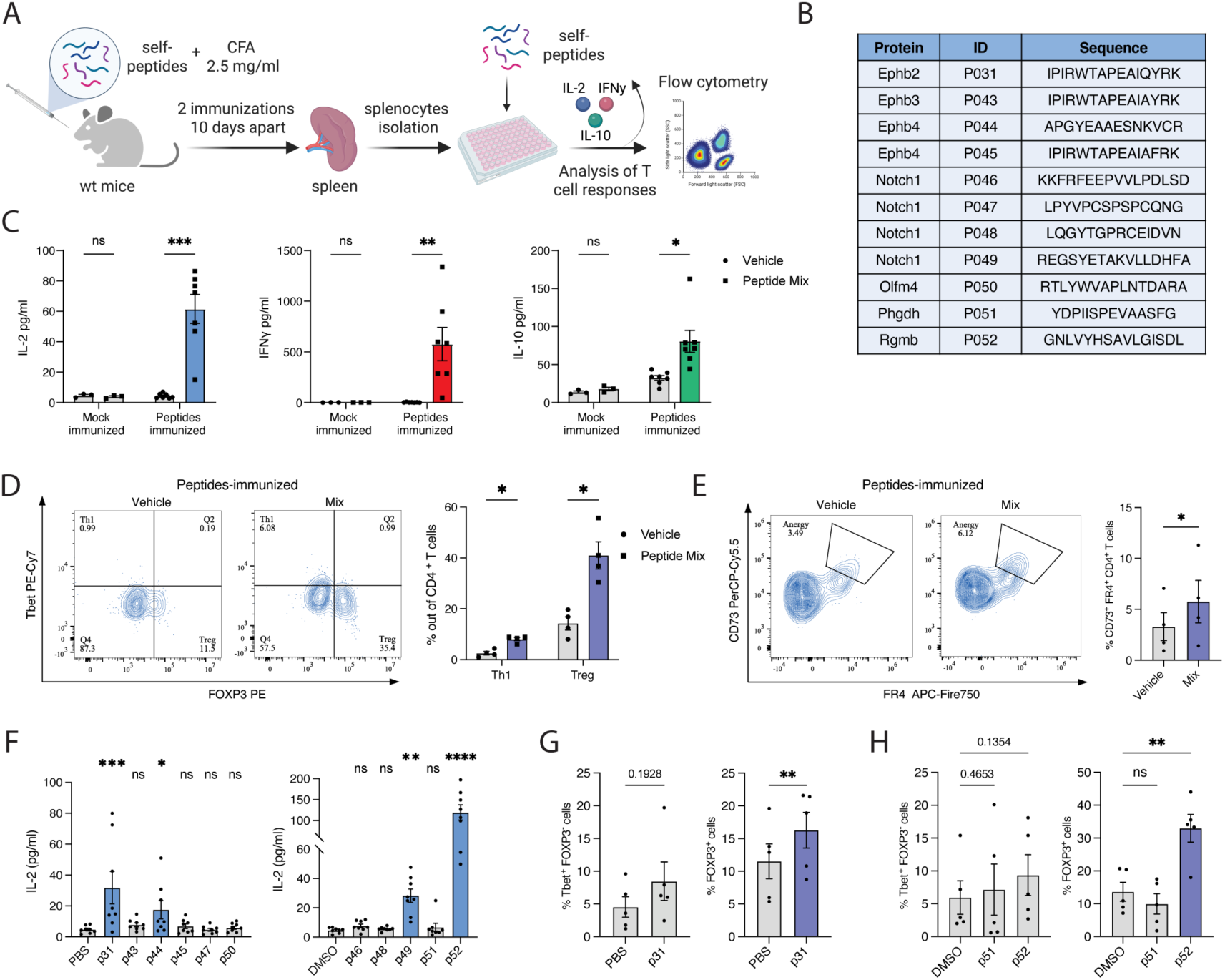
Functional assessment of ISC-specific class II peptides. (**A**) Schematic representation of the *ex vivo* immunogenicity assay. WT mice were immunized with a cocktail of 11 ISC peptides (20 μg each per injection) with Complete Freund’s Adjuvant (CFA). Splenocytes were cultured with a peptide of interest (or a mix of peptides) for 24h. (**B**) List of ISC-specific class II peptides used in the immunogenicity assay. (**C**) Cytokine secretion (IL-2, IFNγ, IL-10) in response to a mix of 11 ISC peptides (2 μg/ml each) by cytometric bead array (CBA) assay. Splenocytes of WT mice immunized with either adjuvant alone or peptides + adjuvant were used. Statistical analysis by multiple paired t-tests with Holm-Šídák correction (α = 0.05), mean ± SEM. \**P* <0.05, \*\**P* <0.01, \*\*\**P* <0.001. (**D**-**E**) CD4^+^ T cell responses upon antigen restimulation. Splenocytes of WT mice immunized with peptides + adjuvant were used. Splenocytes were cultured with a mix of peptides (2 μg/ml each) for 24h. (**D**) Frequency of Th1 (T-bet^+^ FOXP3^-^), Treg (FOXP3^+^) cells out of activated CD4^+^ T cells (CD45^+^ CD3^+^ CD4^+^ CD44^+^) compared to volume-matched vehicle controls. Representative flow cytometry plots are shown. Statistical analysis by multiple paired *t-test*s with Holm-Šídák correction (α = 0.05), mean ± SEM (*n* = 4 mice per group). \**P* < 0.05. (**E**) Frequency of CD73^+^/FR4^+^ cells out of activated non-Treg CD4^+^ T cells (CD45^+^ CD3^+^ CD4^+^ CD44^+^ FOXP3^-^). Representative flow cytometry plots are shown. Statistical analysis by the two-tailed paired *t-test*, mean ± SEM (*n* = 4 mice). \**P* < 0.05. (**F**) IL-2 secretion in response to individual ISC-specific class II peptides (20 μg/ml) by CBA assay. Splenocytes of WT mice immunized with peptides + adjuvant were used. Mice were immunized with a cocktail of 11 ISC peptides. Peptides were split into two groups: PBS-soluble and DMSO-soluble; each group had its volume-matched vehicle control (PBS or DMSO, respectively). Matched (repeated-measures) data were analyzed with a Friedman test, followed by Dunn’s multiple-comparisons test comparing each peptide to the common control (two-sided; PBS- and DMSO-groups were analyzed separately). Reported *P* values are multiplicity-adjusted across all peptide-vs-control tests. Data are mean ± SEM, *n* = 8 mice. \**P* < 0.05, \*\**P* < 0.01, \*\*\**P* < 0.001, \*\*\*\**P* < 0.0001. (**G-H**) Frequency of Th1 (T-bet^+^ FOXP3^-^) or Treg (FOXP3^+^) cells out of activated CD4^+^ T cells (CD45^+^ CD3^+^ CD4^+^ CD44^+^) in response to single-peptide co-cultures compared to volume-matched vehicle-controls. Splenocytes of WT mice immunized with peptides + adjuvant were used. Splenocytes were cultured with a single peptide (20 μg/ml) for 24h. (**G**) Statistical analysis by the two-tailed paired t-tests, mean ± SEM (*n* = 5 mice). \*\**P* <0.01. (**H**) Statistical analysis by multiple paired t-tests with Holm-Šídák correction (α = 0.05), mean ± SEM (*n* = 5 mice). \*\**P* <0.01.

We next sought to identify individual ISC-derived peptides responsible for these effects. Testing each peptide independently revealed four immunodominant ligands derived from Ephb2, Ephb4, Notch1, and Rgmb that elicited significant IL-2 and IFNγ secretion, while Notch1- and Rgmb-derived peptides also induced IL-10 production (**Figures 4F**, **S7C**, and **S7D**). Among these, the Rgmb-derived peptide exhibited the strongest immunomodulatory activity, consistent with the immunomodulating role of Rgmb in intestinal inflammation ^48,49^ (**Figures 4F** and **S7C**). Of note, despite high sequence similarity among ephrin receptor-derived peptides, p31, p43, and p45 (**Figures 4B** and **S5B**), only the Ephb2 peptide (p31) triggered T cell activation, underscoring the fine specificity of self-reactive CD4□ T cells. Importantly, when LP immune cells from immunized mice were restimulated *ex vivo*, responses to these peptides were markedly attenuated, consistent with local peripheral tolerance in the gut microenvironment where these antigens are naturally encountered (**Figure S7E**). As an independent validation of peptide identification, we compared the fragmentation pattern of synthetic Rgmb- and Ephb2-derived peptides with the corresponding peptides detected *in vivo* and observed a strong concordance between the two spectrums, confirming that these immunodominant candidates represent bona fide naturally presented MHC class II ligands (**Figure S7F**).

Finally, stimulation with individual immunodominant peptides selectively expanded FOXP3□ Tregs without increasing Th1 cell frequencies, whereas a non-reactive control peptide had no effect (**Figures 4G** and **4H**). Together, these findings demonstrate that epithelial ISC-derived class II self-peptides are not immunologically inert and can promote regulatory and anergic CD4□ T cell responses (**Figures 4F-4H**, **S7C**, and **S7E**). This establishes a functional link between epithelial antigenic licensing and immune tolerance, setting the stage for examining how inflammatory injury remodels this compartmentalized peptide landscape toward damage-associated antigens during gut inflammation.

### Identification of gut injury class II immunopeptides

To determine how intestinal inflammation reshapes the MHC class II self-peptide landscape, we applied the dextran sodium sulfate (DSS) model of experimental colitis ^18,45^ (**Figure 5A** and **Methods**). Wild-type C57BL/6 mice received 1.5% DSS in drinking water for 7 days, and distal small intestinal tissues were analyzed during the peak of inflammation at day 9 (**Figures S8A** and **S8B**). MHC II immunopeptidomics profiling of villus, crypt, and LP compartments from DSS-treated mice identified more than 13,737 peptides at a 1% FDR, retaining 11,600 sequences after quality control. Restriction to peptides predicted to bind the I-A allele yielded a high-confidence set of 7271 MHC II immunopeptides (**Figures S8C-S8F** and **Table S5**). In marked contrast to steady-state conditions, the villus compartment of DSS-treated mice exhibited an increasing trend in the immunopeptidome diversity, with 3966 peptides and 2039 source proteins identified, indicating a pronounced remodeling of epithelial antigen presentation during inflammation (**Figures S8C** and **S8G**). PCA revealed that gut inflammation disproportionately altered the MHC II ligandome of epithelial compartments relative to immune cells, highlighting epithelial antigen presentation as a sensor of tissue injury (**Figures 5B** and **S8G**). Consistent with this shift, epithelial-enriched class II peptides during ileitis were derived from proteins associated with cellular stress, injury, and immune activation (**Figures 5C** and **S8H**). These included markers of inflammatory cell death pathways, such as the inflammasome-associated protein Gsdmd, the necroptosis regulator Ripk3, and the mitochondrial apoptosis factor Apaf1, as well as immune- and cytokine-related proteins including granzyme A (Gzma), RANK (Tnfrsf11a), interferon signaling components (H2-Aa, Ifngr1), transforming growth factor-β pathway members (Ltbp1, Tgfbr1), and extracellular matrix remodeling proteins such as Loxl1, Mmp2, and Col15a1 (**Figure 5C**). Together, these data indicate a qualitative shift in epithelial MHC II presentation, which alerts epithelial stress, tissue damage, and inflammatory remodeling (**Figures 5C** and **S8H**). Finally, we identified a distinct set of inflammation-specific class II self-peptides detected exclusively during ileitis and absent under steady-state conditions. These 1352 peptides, derived from 589 proteins, included prominent acute-phase response factors such as serum amyloid A1 and A2 (Saa1/2) and Apcs2 (**Figure 5C**). Other acute-phase proteins, lipopolysaccharide-binding protein (Lbp), and the interferon-induced helicase Itih3 were highly enriched in the DSS-treated group (**Figure 5C**). Notably, SAA family members are well-established clinical biomarkers of acute inflammation and, in particular, IBD ^50^. The selective appearance of these acute-phase-associated peptides on MHC II, including epithelial cells, during inflammation suggests that epithelial antigenic licensing is dynamically repurposed under tissue injury to display damage-associated self-antigens, potentially enabling the recruitment and activation of effector CD4 T cells at sites of intestinal inflammation. Together, these data support a model in which epithelial MHC II-dependent licensing is context-dependent, promoting tolerogenic self-antigen display under homeostasis while switching to damage-associated self-antigen signaling during inflammation, thereby positioning the intestinal epithelium as an active sensor and reporter of tissue injury rather than a passive bystander.

**Figure 5.**
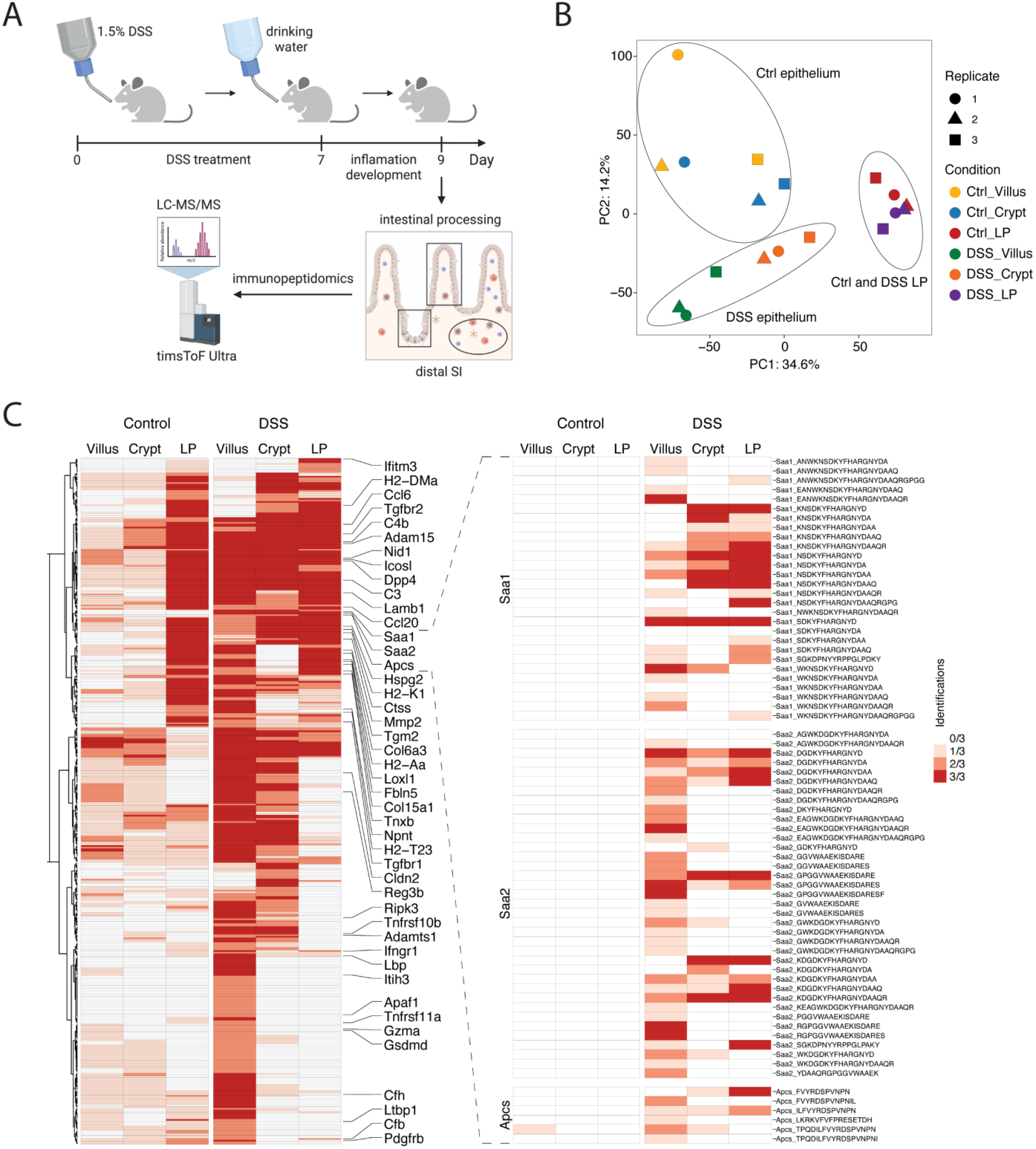
Identification of damage-specific class II immunopeptides. (**A**-**C**) WT mice were treated with 1.5% DSS in drinking water. Villus, crypt, and LP compartments were isolated, and MHC II peptides were profiled (*n* = 3 mice per group). (**A**) Schematic representation of the DSS-induced experimental colitis. (**B**) PCA of immunopeptides from the villus, crypt, and LP compartments in control (Ctrl) vs DSS-treated mice. Axes labeled with variance explained; *n* = 3 mice per condition. (**C**) DSS-enriched source proteins of identified MHC II self-antigens (left panel). Each tile shows the number of biological replicates in which that source protein was detected in the given compartment. DSS exclusive class II immunopeptides of SAA1, SAA2, and Apcs are shown (right panel).

## Discussion

Antigen presentation by MHC II molecules is central to adaptive immune regulation, enabling CD4□ T cells to discriminate between harmless self-antigens and signals that warrant immune activation. In the intestine, an environment continuously exposed to dietary antigens, commensal microbes, and mechanical stress, this balance is particularly critical, as its disruption underlies various gut pathologies, including food allergy, IBD, celiac disease, and colorectal cancer ^14,15,33,51^. While professional APCs, particularly DCs, have long been recognized as key mediators of intestinal tolerance and immunity, recent findings suggest that other APCs may contribute to the complexity of tolerance within the immune compartment ^3,9,34,52–54^. The contribution of IECs to class II antigen presentation has remained controversial, with studies supporting both tolerogenic and pro-inflammatory roles depending on context ^10,14,15,17–19^. In addition, recent studies have shown that epithelial MHC II-expressing cells can induce tolerance to food antigens ^13,55^ or, under infection, orchestrate a compartmentalized immune response ^12^. A strong association between genetic variability at MHC II loci and susceptibility to celiac disease or inflammatory bowel disease (IBD) is well documented ^21,22^. Despite this, the repertoire and nature of peripheral self-antigen presentation by gut APCs, either immune or epithelial cells, remain poorly defined. Advances in mass spectrometry are now enabling us to examine self-immunopeptides associated with tolerance or a specific disease ^23–27^.

Here, we provide a framework to uncover the class II antigenic landscape of the small intestine. Using integrated single-cell transcriptomics, quantitative proteomics, and high-depth *in vivo* gut-immunopeptidomics analysis, we generated the first atlas of MHC II-bound self-peptides across the major compartments of the mucosal layer, the villus, crypt, and LP. We demonstrate that mature enterocytes and ISCs are the dominant epithelial APCs, actively presenting thousands of self-peptides distinct from those displayed by LP immune cells. These epithelial-derived ligands are not randomly sampled but are highly compartmentalized, reflecting epithelial differentiation state, anatomical localization, and physiological function. Malik et al., examined immune responses of intra-epithelial lymphocytes (IELs) and demonstrated their compartmentalized nature, a phenomenon we have now observed in the IEC-ligandome level ^12^. In steady state, the villus MHC II ligandome is enriched for peptides derived from nutrient absorption and metabolic machinery proteins, whereas the crypt ligandome is dominated by ISC-associated proteins, including ephrin signaling components. The close correspondence between transcript abundance, protein expression, and peptide presentation indicates that intestinal APCs broadly sample their steady-state proteome and encode this information directly into the class II ligandome. We propose that this creates a spatial cellular antigenic barcode, allowing CD4 T cells to infer the identity and functional state of the presenting cell and thereby support peripheral tolerance to tissue self. Inducible deletion of MHC II specifically in IECs resulted in a profound collapse of the antigenic compartmentalization, with loss of villus- and crypt-specific self-peptides and compensatory dominance of immune-derived ligands. This collapse was accompanied by ECM remodeling and neutrophil infiltration, demonstrating that epithelial MHC II self-antigens display cellular programs distinct from those of immune cells under steady-state conditions. These findings elevate epithelial MHC II from a passive marker of inflammation ^10,18,21^ to an active organizer of tissue-specific antigen presentation. Functionally, we further demonstrate that epithelial self-peptides are immunologically meaningful. A subset of ISC-derived class II peptides elicited antigen-specific CD4□ T cell responses that were dominated by regulatory T cell (Treg) expansion and anergy-associated programs, even when delivered in a strong immunogenic context. These results provide direct evidence that epithelial self-antigen presentation can bias CD4□ T cell fate toward tolerance and establish a mechanistic link between epithelial antigenic licensing and immune homeostasis. Importantly, our data also reveal that the epithelial MHC II peptidome is dynamic and state-dependent. During DSS-induced ileitis, the epithelial immunoligandome undergoes a marked shift with the emergence of damage-and stress-associated class II self-antigens, including acute-phase proteins and markers of cell death, immune activation, and matrix remodeling. This remodeling was more pronounced in epithelial compartments than in immune cells, suggesting IEC class II ligands are sensors of tissue injury under inflammation. Of note, SAAs protein elevation in gut inflammation was shown to drive pathogenic Th17 cell differentiation, causing additional tissue damage ^56^. In this context, epithelial MHC II licensing appears to be repurposed, from enforcing tolerogenic self-recognition to displaying damage-associated self-antigens. Together, our findings support a model in which epithelial MHC II-dependent licensing operates as a bidirectional communication system between tissue and the immune system. On one hand, epithelial self-peptides are associated with tolerance under homeostasis, while on the other hand, they are associated with damage during inflammation. This framework reconciles prior conflicting observations regarding epithelial MHC II function and extends the classical APC paradigm by positioning IECs, particularly ISCs, as non-canonical APCs that actively encode tissue state into the class II ligandome. Beyond its biological implications, our work establishes immunopeptidomics as a powerful omics-level readout of tissue composition and cellular stress, capable of detecting subtle inflammatory changes and low-abundance immune populations. By defining the epitope-level language through which tissues communicate with CD4□ T cells, this study lays the groundwork for targeted modulation of antigenic landscape as a therapeutic strategy in intestinal inflammation and immune-mediated diseases.

## Acknowledgments

We thank Prof. Ramnik Xavier (Broad Institute) and Prof. Ziv Shulman (WIS) for their critical comments and discussions. We thank Merav Kedmi for help in RNA single-cell experiments and the histology unit. M.B. holds the Ernst and Kaethe Ascher Career Development Chair. This study was supported by research grants from the Center for New Scientists at the Weizmann Institute of Science, the Israel Science Foundation (grant No. 1587/20 and 3775/25), the Helen and Martin Kimmel Institute for Stem Cell Research at The Weizmann Institute of Science, and the Minerva Foundation, with funding from the Federal German Ministry for Education and Research, the Moross Integrated Cancer Center, the Israel Ministry of Science (IMOS, Grant No. 4631), the Dr. Gilbert S. Omenn and Martha A. Darling Weizmann Institute - Schneider Hospital Fund for Clinical Breakthroughs through Scientific Collaborations, a research grant from the Snider Foundation, the Abisch-Frenkel RNA Therapeutics Center, a research grant from the Shimon and Golde Picker, and a research grant from the Herbert K. Bennett Charitable Fund, Dwek Institute for Cancer Therapy Research and a research grant from the Center for New Scientists at the Weizmann Institute of Science.

## Author contributions

V.H. and M.B. conceived the study, designed experiments, and interpreted the results; V.H. and C.W. performed the computational analysis with the assistance of M.C., A.S-P., T.S., B.T.; V.H. carried out all experiments with the help of V.R., M.N., S.L., M.C., P.B., A.H-M., I.D., Z.P., P.Gra.; P.Gre. assisted with immunopeptidomics experiments; C.K. and Y.L. performed MS runs; N.Y., E.T., and Y.S. helped in providing guidance and support; M.B. supervised this study; and V.H. and M.B. wrote the manuscript, with input from all authors.

## Methods

### Mice

All mouse work was performed in accordance with the Institutional Animal Care and Use Committees (IACUC, no. 00330122-3, 07000923-1) of the Weizmann Institute of Science. Mice were housed under specific-pathogen-free conditions at the animal facilities at the Weizmann Institute of Science, Rehovot, Israel. Lgr5-eGFP-IRES-CreERT2 (008875), Villin-CreERT2 (020282), and H2-Ab1^fl/fl^ (037709) mice were purchased from Jackson Laboratory. C57BL/6 wild-type (WT) mice were purchased from either Jackson Laboratory (C57BL/6J; 000664) or Envigo (C57BL/6JOlaHsd). MHC II^ΔIEC^ mice were generated by crossing Villin-CreERT2 and H2-Ab1^fl/fl^ mice. Both male and female age-matched mice aged 10 to 14 weeks were used for all experiments in this study. Littermates of the same genotype, sex, and age were randomly assigned to experimental groups. Cre activity was induced in 10-12 week-old Vil-CreER^T^^2^-H2-Ab1^fl/fl^ mice that express or do not express Cre by intraperitoneal injection (IP) of tamoxifen (Sigma-Aldrich), diluted in corn oil, 2 mg per injection, 5 times every other day. Mice were sacrificed 10 days after the first injection.

### DSS-induced colitis

The Dextran Sulfate Sodium (DSS)-induced colitis model was performed in 10-week-old WT C57BL/6JOlaHsd male mice. Mice received 1.5% DSS (MP Biomedicals LLC) in the drinking water for 7 days, followed by a return to normal drinking water. Mice were sacrificed on day 9 at the peak of inflammation, as determined by weight loss. The distal small intestine was collected and processed for MHC II immunopeptidomics as described below.

### Epithelial cell dissociation, villus and crypt isolation

For all mice, villi and crypts were isolated from the whole small intestine as described before ^14^. Briefly, the small intestine was extracted and rinsed in cold phosphate-buffered saline (PBS). The tissue was opened longitudinally and sliced into small fragments roughly 2 mm long, followed by incubation in 20 mM EDTA-PBS on ice for 75 min. Then, the tissue was shaken vigorously, and the supernatant was collected as fractions. The process was repeated until the supernatant became clear. All the fractions were filtered through a 100 µm pluriSelect filter. For flow cytometry analysis and fluorescence-activated cell sorting (FACS), villi and crypts were dissociated with TrypLE Express (Invitrogen) for 1 min 15 sec at 37°C. The single-cell suspension was then passed through a 40 μm filter and stained for either flow cytometry analysis or FACS sorting (Sony SH800). FACS sorting was performed for bulk RNA-seq, scRNA-seq, and proteomics (see below). For peptidomics experiments, villi and crypts were flash-frozen in liquid nitrogen.

### Immune cell isolation

After villus and crypt isolation, immune cells from the Lamina Propria (LP) were isolated enzymatically by incubating the small intestine with Liberase TM (100 μg/mL, Sigma) and Dnase I (10 μg/mL, Sigma) for 60 min at 37°C while rotating. The tissue was then meshed through the 70 μm filter, and cells were centrifuged at 400g for 5 min. Immune cells were stained for either flow cytometry analysis or FACS. FACS sorting was performed for bulk RNA-seq and proteomics (see below). For peptidomics experiments, Lamina Propria immune cells were flash-frozen in liquid nitrogen. For *ex vivo* peptide restimulation, LP immune cells were resuspended in complete medium (Advanced DMEM, 10% heat-inactivated FBS, 1% GlutaMAX, 10 mM HEPES, 50 µM β-mercaptoethanol, 100 μg/ml Primocin (InvivoGen), counted, and kept on ice until plating.

Spleens and mesenteric lymph nodes (mLN) were processed on ice in washing medium (RPMI-1640, 2% FBS, 25 mM HEPES). Organs were passed through 70 µm strainers; spleens underwent ACK lysis (2 min, RT) followed by quenching in washing medium. Cells were washed (400 × g, 7 min), resuspended in complete medium, counted, and kept on ice until plating.

### Flow cytometry analysis

Single-cell suspensions were prepared as described above. Epithelial and immune cells were stained with EpCAM (Biolegend), CD45 (Biolegend), I-A/I-E (Biolegend), and Dapi-NucBlue (ThermoFisher) for 30’ on ice. Analysis was done on the CytoFLEX S flow cytometer or the Sony SH800 sorter. Further analysis of the data was performed using FlowJo v10.10.1.

### Cell sorting

For the bulk RNA-seq, FACS was performed to sort 10,000 cells into an Eppendorf tube containing 50 μL TCL buffer (QIAGEN) solution with 1% β-mercaptoethanol (Sigma-Aldrich). To isolate MHC II□ and MHC II□ cells, epithelial cells (isolated from villi or crypts) and immune cells of WT mice were stained with the DAPI-NucBlue (ThermoFisher), CD45 (Biolegend, 30-F11), EpCAM (Biolegend, G8.8), I-A/I-E (Biolegend), and gated for MHC II□ or MHC II□ EpCAM^+^ CD45^-^ (epithelial samples) or MHC II□ or MHC II□ CD45^+^ (immune samples) cells. For bulk RNA-seq, the tubes were briefly spun down, immediately frozen on dry ice, and kept at −80°C until ready for RNA isolation. For scRNA-sequencing, the cells were treated as described below. For proteomics, 100,000 MHC II□ EpCAM^+^ CD45^-^ epithelial cells of villi, crypts, and 100,000 MHC II□ CD45^+^ cells were sorted. The tubes were centrifuged at 400g for 5 min, and the supernatant was discarded. The pellets were resuspended in 5% SDS 50mM Tris (pH 7.6) solution, immediately frozen on dry ice, and kept at −80°C until ready for protein isolation.

### Mice immunization

For priming the immune responses against MHC II-associated immunopeptides, a cocktail of 11 commercial synthetic peptides (GenScript; 20 µg of each peptide per mouse) was emulsified in Complete Freund’s Adjuvant (CFA, BD™) at a final *M. tuberculosis* particle concentration of 2.5 mg/ml. A total of 200 µl emulsion per mouse was administered subcutaneously at three sites distributed as follows: two sites in the bilateral scapular region and one in the right dorsolateral rump. For boosting, the same peptide doses were emulsified in Incomplete Freund’s Adjuvant (IFA, BD™) and given 10 days after the prime at the same total volume (200 µl emulsion per mouse) at three sites distributed as follows: two sites in the bilateral lumbar flanks and one in the left dorsolateral rump. Mice were euthanized 10 days after the boost.

Peptide stocks were prepared as follows: PBS-soluble peptides at 2 mg ml□¹ (PBS), and DMSO-soluble peptides at 10 mg ml□¹ (DMSO). Vehicle controls were solvent- and volume-matched to the corresponding experimental setting (control mice for immunization and control wells in *ex vivo* peptide restimulation; see below).

### *Ex vivo* peptide restimulation

For spleen cultures, 1.5 × 10□ cells were plated per well in 100 µl complete medium; for mLN and LP, 0.5 × 10□ cells per well were plated (96-well U-bottom tissue-culture plates). To assess bulk peptide responses, a cocktail of 11 peptides was added to a final concentration of 2 µg ml□¹ per peptide, and the mixture was gently mixed. Control wells received a volume-matched amount of peptide solvents (PBS and DMSO). For individual peptide screening, peptides were added one peptide per well to a final concentration of 20 µg ml□¹ and mixed gently. Vehicle control wells received volume-matched PBS or DMSO, depending on the peptide solvent; for the DMSO-solubilized peptide, the final DMSO concentration in culture was 0.2% (v/v). Cultures were incubated 24 h at 37 °C 5% CO□. After incubation, 50 µl supernatant from each well was harvested for cytokine measurement.

### Cytokine secretion quantification

Cytokines (IL-2, IFN-γ, IL-10) in *ex vivo* co-culture supernatants were quantified using the BD™ Cytometric Bead Array (CBA) Flex Set according to the manufacturer’s instructions, with minor adaptations for low volumes. Briefly, supernatants (50 µl) were incubated with the corresponding capture-bead mixture, followed by PE-conjugated detection reagents, with all incubations performed at room temperature in the dark on a plate shaker. Beads were washed in FACS buffer (PBS, 1% BSA) and acquired on a CytoFLEX S flow cytometer using standard settings. A standard curve was prepared in complete medium in parallel by serial two-fold dilutions covering the assay’s dynamic range. Cytokine concentrations were calculated from median fluorescence intensity (MFI) using a five-parameter logistic fit, applying the same gating template to all samples. For each peptide, cytokine responses were compared against the matched vehicle control (PBS, DMSO, or a volume-matched mix of both) processed in the same plate and run.

### Histochemistry

Tissues were fixed for 16-24 hours in formalin, embedded in paraffin, and cut into 5μm thick sections. Sections were deparaffinized with standard techniques. For H&E staining, slides were stained with Hematoxylin for 1 minute, washed, and then stained with Eosin for 45 seconds.

### Immunofluorescence (IF)

Tissues were fixed for 14 hours in formalin, embedded in paraffin, and cut into 5 μm thick sections. Sections were deparaffinized with standard techniques, incubated with primary antibodies for Ly6G (1:100, Biolegend 127606), PDPN (1:100, BioLegend 127402), Olfm4 (1:200) (Cell Signaling Technology, 39141S), Mmp9 (1:100, HUA-ET1704-69-HuaBio) and E-cadherin (1:100) (BD, BD610182) overnight at 4°C, followed by secondary antibodies incubation (1:400, Abcam) at room temperature for 30 min and Hoechst 33342 (1:1000) (TargetMol, T5840). Slides were mounted with Fluoromount-G (SouthernBiotech) and sealed.

### Bulk population RNA purification and cDNA preparation

Libraries were prepared using a modified SMART-Seq2 protocol^102^. RNA lysate cleanup was performed using RNAClean XP beads (Agencourt), followed by reverse transcription with Maxima Reverse Transcriptase (Life Technologies) and whole transcription amplification (WTA) with KAPA HotStart HIFI 2 3 ReadyMix (Kapa Biosystems) for 18 cycles. WTA products were purified with Ampure XP beads (Beckman Coulter), quantified with Qubit dsDNA HS Assay Kit (ThermoFisher), and assessed with a high-sensitivity DNA chip (Agilent). RNA-seq libraries were constructed from purified WTA products using the Nextera XT DNA Library Preparation Kit (Illumina). The libraries were sequenced on an Illumina NovaSeq 6000 using the SP kit.

### Hash-tagged single-cell RNA-sequencing

Single-cell RNA-sequencing (scRNA-seq) libraries were prepared using the Chromium single-cell RNA-seq platform (10x Genomics). Isolated crypt epithelial cells were sorted into a cooled 1.7 mL tube with 0.04% BSA in PBS using a Sony SH800 cell sorter. Cells for each mouse were stained using Biotin anti-mouse CD326 (EpCAM) (cat #118204) and CD45 (Biolegend, 30-F11) antibodies, followed by staining using TotalSeq™ PE Streptavidin (B0951 - B0955). 20,000 EpCAM^+^ CD45^-^ cells were sorted for each mouse and then pooled according to the manufacturer’s instructions. 30,000 single-cell suspension was loaded onto a Next GEM Chip G targeting 15,000 cells, then run on a Chromium Controller instrument to generate a GEM emulsion. Single-cell 3’ RNA-seq libraries and cell surface protein libraries were generated according to the manufacturer’s protocol (Chromium Single Cell 3’ Reagent Kits User Guide (v3.1 Chemistry Dual Index). Final libraries were quantified using NEBNext Library Quant Kit for Illumina (NEB) and high-sensitivity D5000/D1000 TapeStation (Agilent). Libraries were pooled based on the targeted cell number, aiming for ∼20,000 reads per cell for gene expression libraries and ∼5,000 reads per cell for cell surface protein libraries. Pooled libraries were sequenced on a NovaSeq 6000 instrument using an S1 100 cycles reagent kit (Illumina).

### Proteomics workflow

#### Sample preparation

Cells were lysed with 5% SDS in 50 mM Tris-HCl. Lysates were incubated at 96 °C for 5 min, followed by six cycles of 30 s of sonication (Bioruptor Pico, Diagenode, USA). Proteins were reduced with 5 mM dithiothreitol and alkylated with 10 mM iodoacetamide in the dark. Phosphoric acid was added to the lysates to a final concentration of 1.2%. 90:10% methanol/50 mM ammonium bicarbonate was then added to the samples. Each sample was then loaded onto S-Trap microcolumns (Protifi, USA). Samples were then digested with trypsin for 1.5 h at 47 °C. The digested peptides were eluted using 50 mM ammonium bicarbonate; trypsin was added to this fraction and incubated overnight at 37 °C. Two more elutions were made using 0.2% formic acid and 0.2% formic acid in 50% acetonitrile. The three elutions were pooled and vacuum centrifuged to dry. Samples were kept at −80 °C until analysis.

#### Liquid chromatography

Each sample was loaded using split-less nano-Ultra Performance Liquid Chromatography (nanoElute2, Bruker, Germany). The mobile phase was: A) H2O + 0.1% formic acid and B) acetonitrile + 0.1% formic acid. The peptides were separated using an Aurora column (75μm ID x 25cm, IonOpticks) at 0.3 µL/min. The column was kept at 50 °C. Peptides were eluted from the column into the mass spectrometer using the following gradient: 2% to 35%B in 60 min, 35% to 95%B in 0.5 min, maintained at 95% for 6.77 min.

#### Mass spectrometry

The column was connected to the CaptiveSpray electrospray ionization source to the timsTOF Pro mass spectrometer (Bruker, Germany). The MS was operated in Data Independent Acquisition Parallel Accumulation Serial Fragmentation (DIA-PASEF) mode. MS1 mass range was 100-1,700 m/z. TIMS ramp was set to 100msec with a range of 0.6-1.6 1/K^0^. DIA windows were set to 25 Th with 1 Th overlap, range of 300 to 1,189 m/z with a cycle time of 1.48 sec.

### MHC II immunopeptidomics workflow

#### Immunopeptidome sample processing

MHC II purification was performed according to published procedures, with study-specific modifications. Villus and crypt samples of ∼500mg and immune samples of 2×10^7^ cells, representing a whole small-intestinal isolate, were resuspended in a lysis buffer (containing 0.25% sodium deoxycholate, 0.2mM iodoacetamide, 1mM EDTA, protease inhibitor cocktail (SigmaAldrich), 1mM PMSF, and 1% octyl-β-D-glucopyranoside in PBS) and subject to two rounds of sonication at 30% amplitude for 15 sec. Lysates were kept on rotation at 4°C for 1 hour and then cleared by two consecutive centrifugations at 4°C, 48,000g for 40 and 20 minutes, respectively. The supernatant was incubated in a pre-clearing column containing Protein G resin (GenScript) for 20 min at 4°C on rotation and passed through it. Cleared lysates were incubated for 1.5 h at 4 °C under rotation with M5/114.15.2 anti-MHC II mAb (Bio X Cell) covalently coupled to Protein G resin (GenScript). The same Protein G resin was used for the pre-clearing step. Immunoaffinity columns were washed first with one column volume of lysis buffer, four column volumes of 400 mM NaCl, 20 mM Tris-HCl (pH 8.0), and finally with two column volumes of 20 mM Tris-HCl (pH 8.0). MHC II-peptide complexes were eluted at room temperature by adding 330 μL of 1% trifluoracetic acid (TFA) in a total of 3 elutions per sample. Eluted MHC II peptides and the MHC II complexes were loaded into the wells of a Sep-Pak tC18 100mg Sorbent 96-well plate (Waters), which were prewashed with 80% acetonitrile (ACN) in 0.1% TFA and with 0.1% TFA. The peptides were separated from the much more hydrophobic MHC II molecules by eluting them with 31% ACN in 0.1% TFA.

#### LC-MS/MS analysis of MHC II peptides

The immunopeptides were dried by vacuum centrifugation, resolubilized with 0.1% TFA and 5mM TCEP in water, and separated using reversed-phase chromatography using the nanoElute2 system (Bruker), with an Aurora analytical column, 0.075 × 250 mm (IonOpticks, Australia). Mobile phase A: H2O+0.1% formic acid, B: acetonitrile+0.1% formic acid. The chromatography system was coupled by electrospray to tandem mass spectrometry to the timsTOF Ultra (Bruker). Peptides were eluted with a linear gradient from 5 to 23% B in 37 minutes, 23 to 35% B in 8 minutes, then 4 minutes to 95% at a flow rate of 0.25 μL/min. Data were acquired using a data-dependent parallel accumulation-serial fragmentation (DDA-PASEF) method. Full-scan MS spectra were acquired at a m/z, 100-1700 mass range. Mobility range of 0.64 to 1.7 1/K0, 100msec ramp time. MS/MS was acquired at a mass range of 100-1,700 m/z, 5 PASEF ramps per cycle, triggering on charges 1 to 4. Target intensity of 20,000, intensity threshold of 500. Collision energy was set to 20 eV at 0.7 1/K0 up to 70eV at 1.68 1/K0.

## Computational methods

### Bulk RNA-sequencing data analysis

#### Read QC, trimming, and alignment/quantification

Samples were demultiplexed using bcl2fastq v2.20. Adapter and quality trimming were performed with cutadapt v2.7 (minimum length 25 nt, quality cutoff Q10). Read quality was assessed with FastQC v0.11.8. Trimmed reads were aligned to the Mus musculus reference genome GRCm39 with STAR v2.7.3a (with EndToEnd option) against the Ensembl release 105 gene annotation (GTF). Gene-level counts were obtained with HTSeq-count v0.12.4. Only uniquely mapped reads were used to determine the number of reads that map to each gene (intersection-strict mode).

#### Normalization and differential expression

Genes with fewer than 5 counts in all the samples were filtered. Differential expression was performed in R v4.2.2 using DESeq2 v1.38.3 ^59^ with the betaPrior, cooksCutoff, and independentFiltering parameters set to False. Raw P values were adjusted for multiple testing using the procedure of Benjamini and Hochberg. Differentially expressed genes were determined by a p-adj of <0.05 and absolute fold changes >1.5 and a count of at least 30 in at least one sample. Heatmap plotting was done with the ComplexHeatmap R package v2.14, using scaled log2 DESeq2-normalized counts for each gene. For gene-signature scoring, epithelial marker gene sets were curated from our previously published epithelial dataset of the small intestine, with each epithelial subtype treated as a separate gene set. For villus and crypt populations, raw DESeq2 counts were ranked per sample using rankGenes, and the singscore R package v1.18.0 ^65^ was applied using the simpleScore function with default random background sampling (equal size). The resulting per-sample signature score was used for downstream visualization. Specifically, for each epithelial gene set, log□ fold changes between MHC II□ and MHC II□ samples were computed, and significance was assessed by two-sample t-tests with Benjamini-Hochberg adjustment. Gene Ontology (GO) enrichment was performed in R (clusterProfiler package v4.8.3, enrichGO function) using the *Mus musculus* annotation (org.Mm.eg.db), testing Biological Process terms separately for up- and downregulated gene sets; significance was defined as FDR < 0.05.

#### Deconvolution (CDSeqR)

Cell composition was inferred from bulk gene-count data using the CDSeqR v1.0.8. R package ^57^ in complete (reference-free) deconvolution mode, separately for epithelial (villus and crypt) and immune samples. The number of latent cell types, K=15, was chosen after manual inspection for both epithelial and immune samples. CDSeqR returned sample-level cell-type proportion estimates and cell-type-specific expression profiles. For biological interpretation, components were annotated against published scRNA-seq datasets of small intestinal epithelial and immune cells. The acquired proportions were averaged per condition (MHC II□ vs MHC II□ samples of villus, crypt, and immune cohorts) for visualization.

### scRNA-sequencing data analysis

#### scRNA-seq pre-processing and demultiplexing

The raw sequencing data were processed using the Cell Ranger toolkit (version 7.0, 10x Genomics). FASTQ files were generated from Illumina sequencer output using the bcl2fastq software (version 2.20.0.422) with default parameters for demultiplexing and converting BCL files to FASTQ format. The FASTQ sequences were then aligned to the mouse reference genome (GRCm38-mm10) using the STAR aligner embedded within Cell Ranger.

#### Quality Control of Single-Cell RNA Sequencing Data

The raw gene expression matrices were imported into R (version 4.1.1) and analyzed using the Seurat R package (version 5.0.1). The dataset included a total of 16,451 isolated from 5 individual mice. A stringent quality control process was employed to remove low-quality cells, which could confound downstream analyses. Cells were filtered based on the following criteria: (i) cells expressing fewer than 300 or more than 6000 genes were excluded to remove potential empty droplets or doublets; (ii) cells with a total UMI (Unique Molecular Identifier) count exceeding 35,000 were discarded to avoid potential doublets; and (iii) cells with more than 15% of reads mapping to mitochondrial genes were excluded to eliminate cells likely undergoing apoptosis or other stress-related processes. This filtering process ensured that only high-quality, biologically relevant cells were retained for subsequent analysis.

#### Normalization, Scaling, and Dimensionality Reduction

Normalization and scaling of the raw gene expression data were performed using the Seurat function SCTransform, which applies a regularized negative binomial regression to model and remove technical noise while retaining biological variability. After normalization, dimensionality reduction was carried out using Principal Component Analysis (PCA) on the SCT-normalized data, implemented via the Seurat RunPCA function. The top 11 principal components, representing the most significant sources of variation in the data, were selected for downstream analyses. For cell clustering, a k-nearest neighbors (k-NN) graph was constructed using the Seurat functions FindNeighbors and FindClusters, facilitating the identification of distinct cell populations. Finally, two-dimensional visualization of the data was achieved by applying Uniform Manifold Approximation and Projection (UMAP) using the RunUMAP function in Seurat, with the top 11 principal components, n_neighbors value of 30, and min_dist value of 0.3 serving as input. This approach identified 31 transcriptionally distinct clusters.

#### Identification and Annotation of Cell Clusters

To identify differentially expressed genes (DEGs) within each cell cluster, the FindAllMarkers function in Seurat was utilized. This function performs a pairwise comparison between each cluster and all other clusters, enabling the identification of genes that are significantly upregulated or downregulated in specific clusters. The top differentially expressed genes were selected based on their statistical significance and log-fold change, providing a basis for subsequent cluster annotation.

Cell clusters were annotated by integrating the results of the DEGs analysis with multiple identification strategies to ensure accurate classification of cell types and states. These strategies included:

1. **Canonical Cell Type-Specific Markers**: Well-established markers specific to certain cell types were used to identify and annotate the major cell populations within each cluster.
2. **Biological State-Related Markers**: Markers associated with specific biological processes, such as stemness, proliferation, or differentiation, were employed to refine the annotation further, particularly for identifying dynamic states within cell populations.

To further characterize the cell cycle dynamics within the identified clusters, the CellCycleScoring function was applied. This function calculates the average expression levels of genes associated with three distinct phases of the cell cycle: S phase (DNA synthesis), G1 phase (cell growth before DNA replication), and G2M phase (cell growth before mitosis). By scoring cells based on their expression of these phase-specific markers, we were able to infer the proliferative state of cells within each cluster, providing additional insights into their biological roles. 31 initial clusters were consolidated into 14 meta-clusters based on their DEGs and cell-cycle dynamics, yielding major epithelial subtypes and their differentiation states.

### MHC II machinery and lipid metabolism signature analysis

To quantify the expression of the MHC II machinery and lipid metabolism signatures within our single-cell RNA sequencing (scRNA-seq) dataset, we employed the UCell R package ^61^ (https://github.com/carmonalab/UCell). UCell is a computational tool specifically designed for scoring gene signatures across scRNA-seq data, enabling the identification and comparison of cellular phenotypes based on predefined gene sets.

The MHC II machinery signature included the following genes to represent antigen loading and presentation: *H2-Aa*, *H2-Ab1*, *Cd74*, *H2-Eb1*, *H2-DMa*, *H2-DMb1, Ciita*. The genes for lipid metabolism were selected to represent the top GO terms of MHC II^+^ villus involved in lipid digestion, transport, and synthesis (**Figure S1G**): *Apoa1*, *Apoa4*, *Apoc3*, *Apoe*, *Cidec*, *Gpat3*, *Dgat2*, *Lipa*, *Slc27a2*, *Slc27a4*, *Scarb1*, *Cd36*, *Acox2*, *Acox3*, *Fabp2*, *Asah2*.

UCell calculates an enrichment score for the gene signature within each cell, allowing for the identification of cells that express the signature at varying levels. The analysis was conducted according to the standard workflow described in the UCell documentation.

### Mass-spectrometry data analysis

#### Proteomics raw data processing

Raw mass spectrometry data were processed with Spectronaut (Biognosys, v19.1) using the default directDIA+ workflow. Searches were performed against the Mus musculus (mouse) UniProtKB/SwissProt reference proteome (version 01-2025). Enzyme: Trypsin/P; missed cleavages: up to 2. Fixed modification: carbamidomethyl-C; variable modifications: oxidation-M and protein N-terminal acetylation. FDR was controlled at 1% at precursor and protein levels using Spectronaut’s q-value. Cross-run normalization was enabled. Quantification was based on unique peptides. Other parameters followed Spectronaut defaults.

#### Proteomics downstream analysis

8,140 proteins that have >60% valid values (in at least 2 replicates) in at least one group were identified. 7,730 proteins were represented by at least two detected peptides. Differential expression analysis was performed using the ‘Limma’ package v007 in R, after logarithmic transformation. Imputation was implemented to replace missing values. Differentially expressed (DE) proteins were determined by a cutoff of: absolute fold change>2, p-value <0.05, and at least 2 peptides detected by mass-spectrometry.

Unsupervised analysis was performed to identify patterns in protein expression by clustering 5025 proteins (**Figure S2D**). These proteins were identified as DE in at least one pairwise comparison. K-Means clustering was performed. Standardized, log2-normalized intensities were used for the clustering analysis. Clustering analysis was performed with RStudio v4.2.2 (RStudio Team, 2020).

Proteomics and bulk RNA-seq datasets were integrated for downstream analysis using the 7,622 genes detected in both experiments. Spearman’s rank correlation coefficient (ρ) and associated p-values were computed between log□ fold-change values with RStudio v4.2.2 (RStudio Team, 2020). Only genes classified as DE in at least one dataset were considered. DE status was determined according to the platform-specific criteria applied in each analysis.

### MHC II immunopeptidomics raw data analysis

The raw mass-spectrometry (MS) data files were analyzed using the FragPipe computational proteomics platform version 23.1 with the default immunopeptidomics workflow ^66^. MSFragger v4.3, a search engine incorporated in the FragPipe framework, was used to search the peak lists against the Ensembl v113 Mus musculus (mouse) database and a file of frequently observed contaminants.

N-terminal acetylation (42.010565 Da) and methionine oxidation (15.994915 Da) were set as variable modifications. The enzyme specificity was set as unspecific. A decoy dataset was generated using a “random” setting. A false discovery rate (FDR) of 0.01 was applied for peptide spectrum match (PSM), and no protein FDR was set. Peptide intensities were extracted using the IonQuant method, embedded in FragPipe. The ‘match between runs’ and normalization options were enabled. Two independent LC–MS/MS immunopeptidomics acquisitions were performed (“Run 1” and “Run 2”; **Table S4**) to increase confidence in the identified peptide repertoire. For QC and qualitative summaries, identifications from both runs were pooled and analyzed together. For quantitative analysis (principal component analysis (PCA)), only Run 1 was used.

NetMHCIIpan v.4.3 ^36^ was utilized to predict peptide binding affinity (binder <10% rank). “Unique” peptides were defined as non-redundant sequences of 9 to 25 amino acids after removal of frequently observed contaminants. For all immunopeptidomics QC figures and downstream analysis (except the percentage of binding sequences plots), only peptides classified as binders were used. Peptide cores were defined by NetMHCIIpan v.4.3.

### Gene Ontology (GO) enrichment analysis

Gene Ontology (GO) enrichment analysis was performed on gene sets derived from either (i) compartment-enriched peptide cores (intestinal immunopeptidome under homeostasis) or (ii) inflammation-enriched source proteins per compartment (intestinal immunopeptidome under inflammation). A peptide core was defined as enriched for a given compartment if its corresponding peptide(s) were detected in ≥2 biological replicates of the target compartment and in ≤1 replicate across all non-target compartments. Immunopeptide source proteins were classified as inflammation-enriched within a given compartment if at least one corresponding peptide was detected in ≥2 biological replicates under inflammation and in ≤1 biological replicate under control for that same compartment. Enriched GO terms were identified using the R package clusterProfiler v4.8.3 ^60^ with “GO Biological Process” and gene annotation provided by the org.Mm.eg.db database. Pathways with adjusted P values below 0.05 were considered (Benjamini-Hochberg FDR).

### Principal Component Analysis

MS data was analyzed using the R package DEP. The FragPipe output table (combined_peptide.tsv) was converted to MaxQuant (MQ)-like output (peptides.txt) using the custom immunopeptidomics analysis pipeline as previously described (https://github.com/YSamuelsLab/MetaPept2) ^67^. The converted table was used as input for DEP, and only peptides classified as binders (NetMHCIIpan v.4.3 <10% rank) were kept. Missing values were handled by filtering with thr = 0, followed by variance-stabilizing normalization (VSN) and left-censored imputation using MinProb (q = 0.01). Principal component analysis (PCA) was performed on normalized, imputed peptide intensities (top 3000 most variable peptides).

### Retention time and hydrophobicity index

Hydrophobicity values for total PSM peptides were predicted with the R package protViz version 0.7.7 ^68^ and were plotted against the experimental retention time. Applied linear regressions were built with the statistics library from the scipy Python package version 1.8.0 ^69^.

### Spectra validation

Light synthetic peptides for spectra validation were ordered from GenScript, as HPLC grade (≥85% purity). These were analyzed using the same LC-MS/MS system and acquisition parameters as indicated above for the endogenous peptides. The data were processed with MQ v.2.1.3.0 using the following parameters: all FDRs were set to 1, and the individual peptide mass tolerance was set to false. The intestinal immunopeptidomics data were reprocessed in MQ following the same parameters. MQ spectra from endogenous and synthetic runs were correlated in Prosit (termed mirror plots) ^70^.

### Statistical analysis

Statistical analysis was performed using Prism (GraphPad) and R v4.3.1. The specific tests applied are detailed in the corresponding figure legends and computational methods.

## Supplemental figures

**Figure S1.**
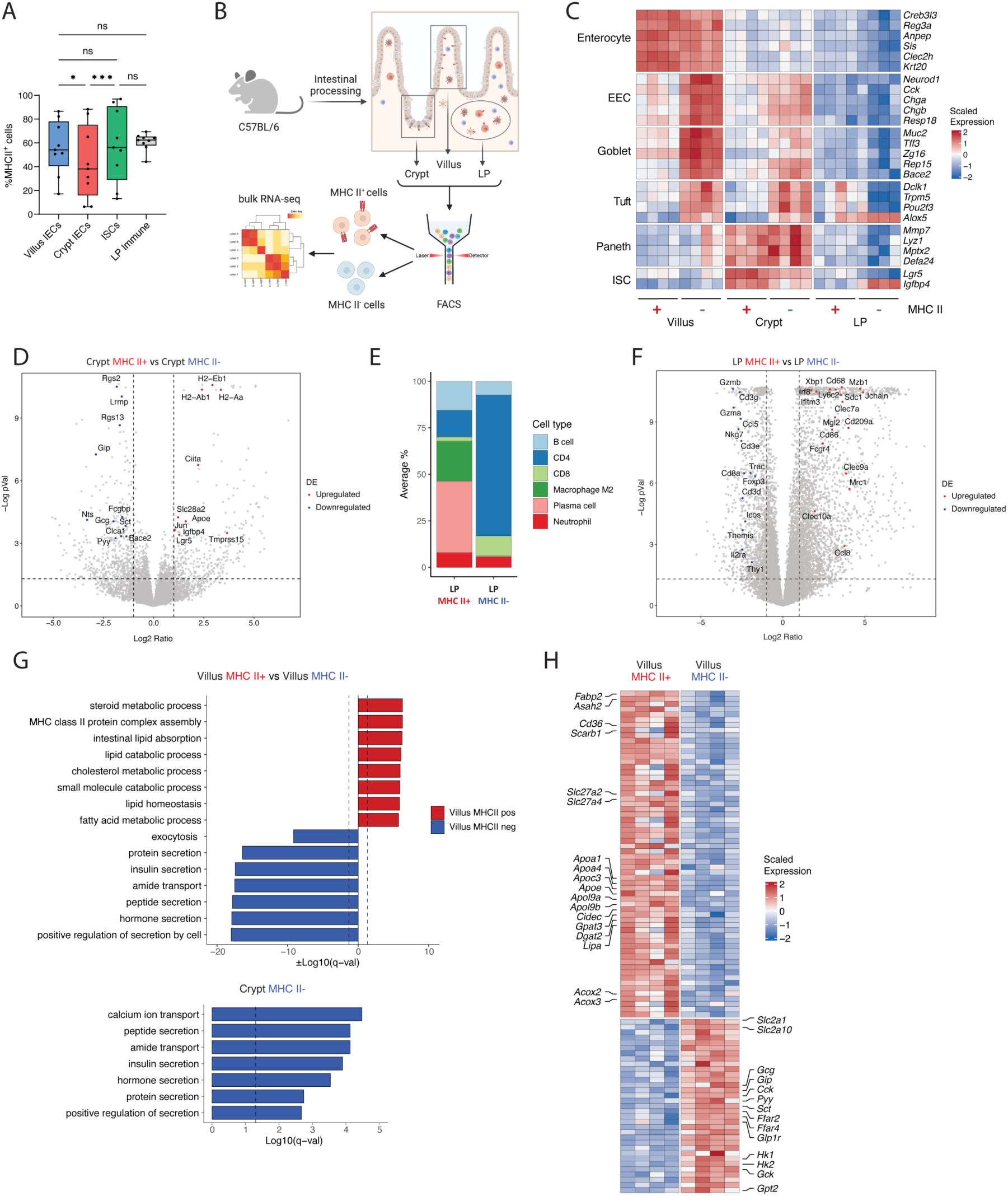
Quality control of bulk RNA-sequencing. **(A)** Frequency of MHC II⁺ EpCAM^+^ CD45^-^ IECs in villus and crypt compartments, MHC II⁺ EpCAM^+^ CD45^-^ eGFP^high^ ISCs in crypt, and MHC II⁺ CD45^+^ immune cells in LP measured by flow cytometry. Frequency is represented as box-and-whisker plot (center line = median; box = 25th-75th percentiles; whiskers = 10th-90th percentiles); *n* = 9 mice. Significance was established by repeated-measures one-way ANOVA with Geisser-Greenhouse correction, followed by Šídák-adjusted two-sided post-hoc tests. \**P* < 0.05, \*\*\**P* < 0.001. **(B to H)** Bulk RNA-seq of MHC II⁺ and MHC II⁻ IECs from villus and crypt and immune cells from LP of WT mice (n = 4). **(B)** Schematic representation of cell sorting strategy for Bulk RNA-seq. (**C)** Expression of IECs markers across the villus, crypt, and LP compartments. Tiles show scaled expression (row-wise z-score) per sample; rows are grouped by lineage and manually ordered; columns are manually ordered by compartment and MHC II status (+/− labels on the x-axis). Marker selection: enteroendocrine, goblet, and tuft genes are differentially expressed (DE) between MHCII⁺ and MHCII⁻ samples within both villus and crypt; stem genes are DE between MHCII⁺ and MHCII⁻ in crypt; enterocyte and Paneth markers are DE between villus and crypt IECs (MHCII⁺ and MHCII⁻ samples combined). DE criteria are provided in **Methods**. **(D)** Volcano plot of DE genes between MHC II⁺ and MHC II⁻ crypt IECs. The x-axis shows log₂ fold change (MHC II⁺/MHC II⁻), with positive values indicating genes upregulated in MHC II⁺ samples and negative values indicating genes upregulated in MHC II^-^ samples; the y-axis shows −log₁₀ of the Benjamini-Hochberg-adjusted *P-value* calculated by DESeq2 ^59^ (**Methods**). Dashed lines indicate DE thresholds: |log₂FC| = 1 (vertical) and padj = 0.05 (horizontal). Highlighted genes: red, MCHII^+^ upregulated; blue, MCHII^-^ upregulated; **(E)** Averaged immune cell-type proportions inferred by CDSeqR ^57^ and annotated using previously published data ^14^. **(F)** Volcano plot of DE genes between MHC II⁺ and MHC II⁻ immune cells. Axes and threshold are as in (**D**). Highlighted genes: red, MCHII^+^ upregulated; blue, MCHII^-^ upregulated. **(G)** Gene Ontology (GO) of biological process enrichment for DE genes between MHC II⁺ and MHC II⁻ IECs within villus and crypt compartments. Upregulated and downregulated gene sets were analyzed separately using clusterProfiler R package ^60^ (**Methods**). Bars represent ±log₁₀ (q-value); red color indicates enrichment in MHC II⁺ cells, and blue indicates enrichment in MHC II⁻ cells; dashed line marks *q* = 0.05. **(H)** Top 100 DE metabolic genes between MHC II⁺ and MHC II⁻ villus IECs. Tiles show row-z-scored log₂-normalized expression per sample; rows are unclustered; columns are ordered by MHC II status. DE was determined using DESeq2 ^59^ (Benjamini-Hochberg padj < 0.05; |log₂FC| > 1). The top 100 genes ranked by |log₂FC| were annotated to metabolic processes and plotted. Representative genes are labeled.

**Figure S2.**
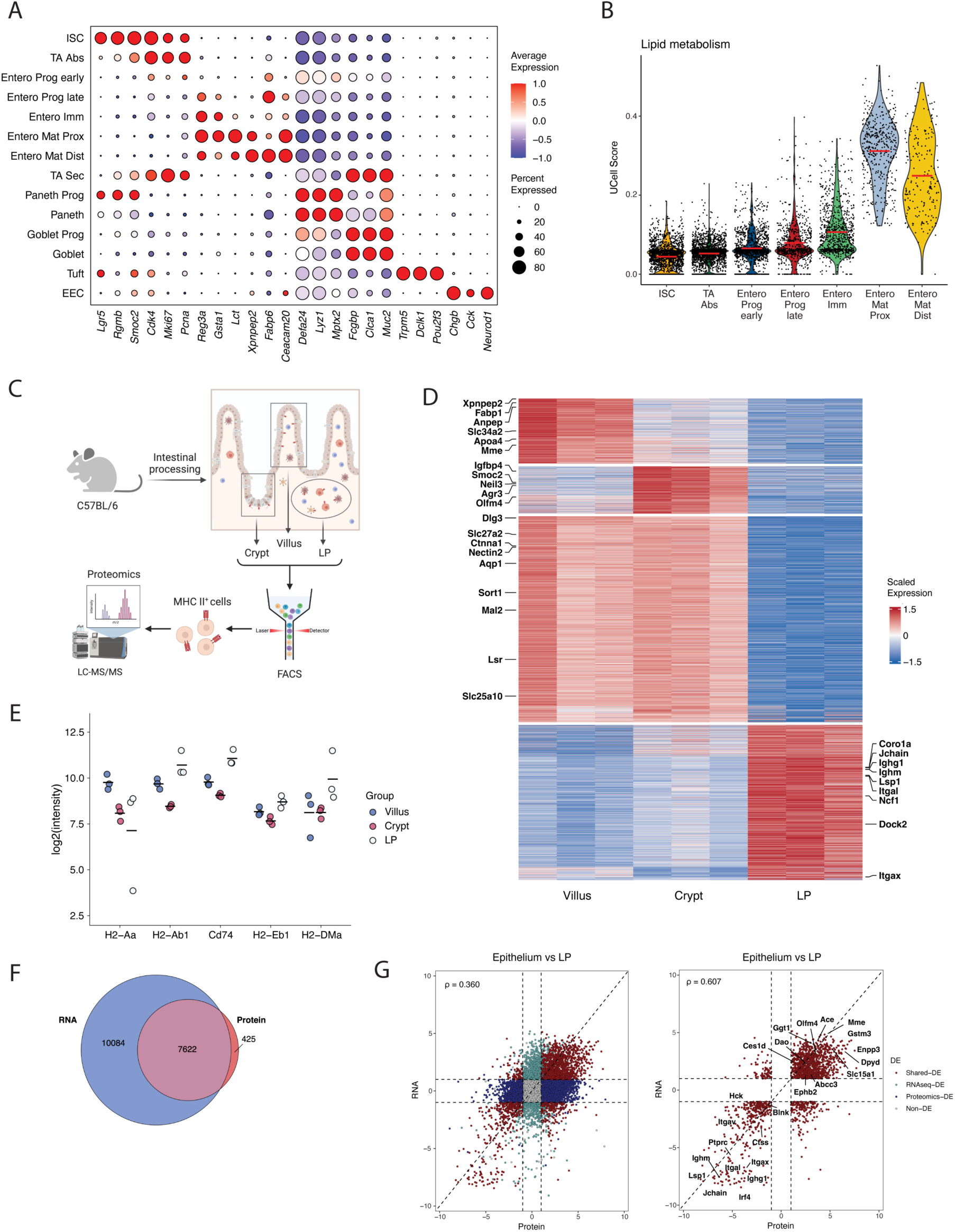
Quality control of scRNA-sequencing and proteomics. **(A)** Dot plot showing expression of cell-subset markers in small-intestinal epithelial scRNA-seq. Rows show annotated epithelial cell subsets, and columns show marker genes. Dot size indicates the fraction of cells in each cluster with detectable expression (non-zero counts) and dot color indicates mean expression per cluster, z-scored per gene across clusters. **(B)** UCell scores ^61^ (**Methods**) for the lipid-metabolism gene signature across annotated epithelial cell types. Each dot represents a single cell; red lines indicate the median score per cluster. **(C)** Schematic of cell sorting strategy for proteomics. **(D)** Proteome heatmap of cell-type-specific protein markers in MHC II⁺ villus, crypt, and LP immune cells. Tiles show row-z-scored log₂ protein abundance per sample; columns are grouped by compartment, and rows are ordered by unsupervised analysis (**Methods**). Row labels indicate representative protein markers. **(E)** MHC II machinery-related protein expression in MHC II^+^ villus and crypt IECs, and LP immune cells. **(F)** Area-proportional Venn diagram of genes detected by bulk RNA-seq or proteins detected by LC-MS/MS proteomics across all MHC II⁺ cells. Numbers indicate feature counts. **(G)** Scatter plots comparing log₂ fold change at the protein (x-axis) and RNA (y-axis) levels for MHC II⁺ populations between epithelial (villus and crypt IECs samples combined) and LP immune cells. Dashed lines mark |log₂FC| = 1. Spearman’s ρ correlation is reported for all DE genes/proteins (Spearman’s ρ = 0.360, left panel) and for only shared DE genes/proteins (Spearman’s ρ = 0.607, right panel).

**Figure S3.**
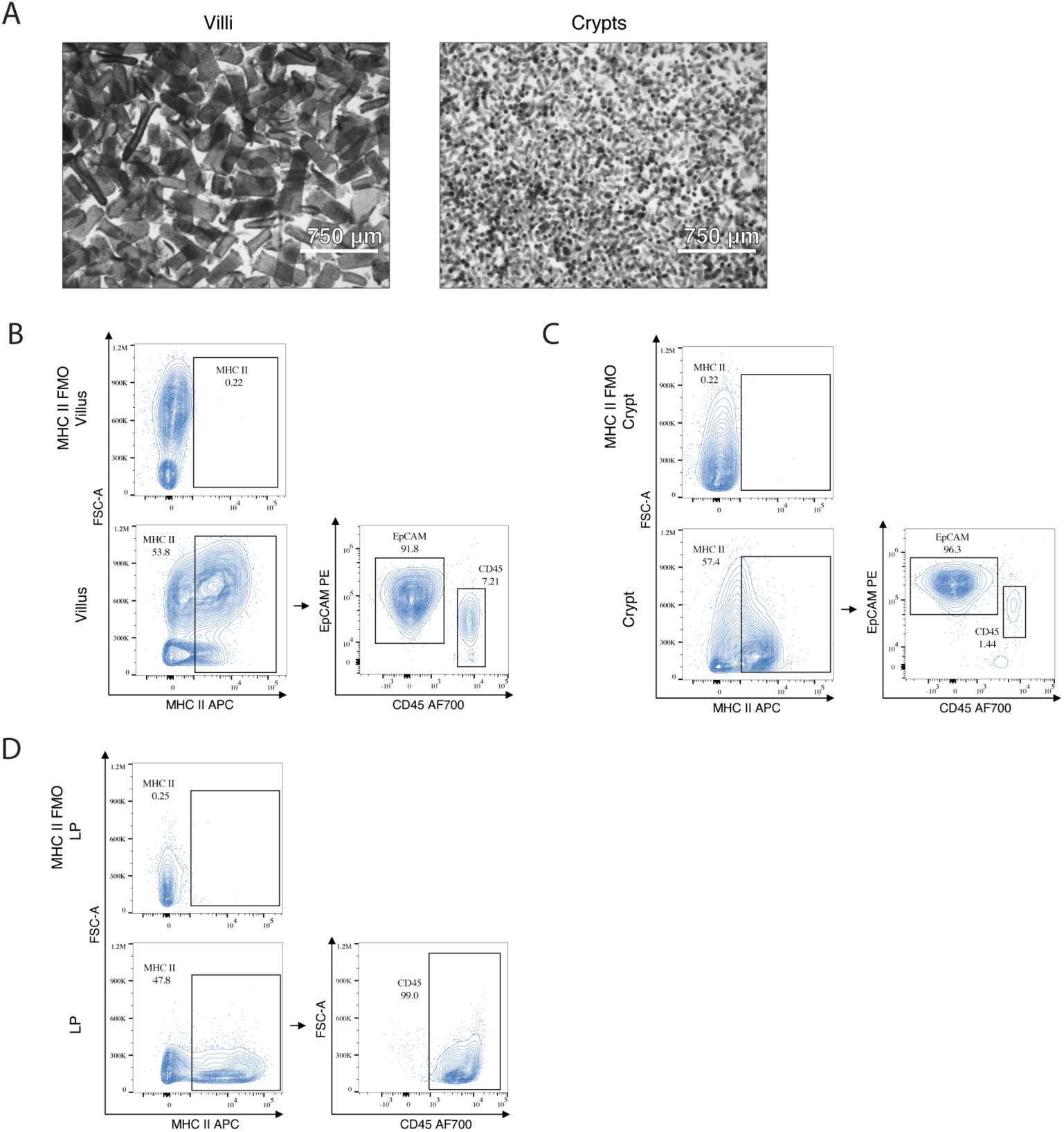
Purity of intestinal compartment samples profiled by immunopeptidomics. **(A)** Representative bright-field micrographs of isolated villus (left) and crypt (right) fractions. Scale bar, 750μm. **(B-D)** Flow-cytometry quality control of intestinal compartment samples used for MHC II peptidomics. Percentages indicate the fraction of cells (CD45^+^ or EpCAM^+^) within the current gate. Representative plots are shown for villus **(B)**, crypt **(C)**, and LP **(D)**; the upper plots display the Fluorescence Minus One (FMO) control for MHC II stain. Sample purity was assessed by the proportion of EpCAM^+^ CD45^-^ cells for the villus and crypt fraction, or CD45⁺ LP cells out of live single MHC II⁺ cells.

**Figure S4.**
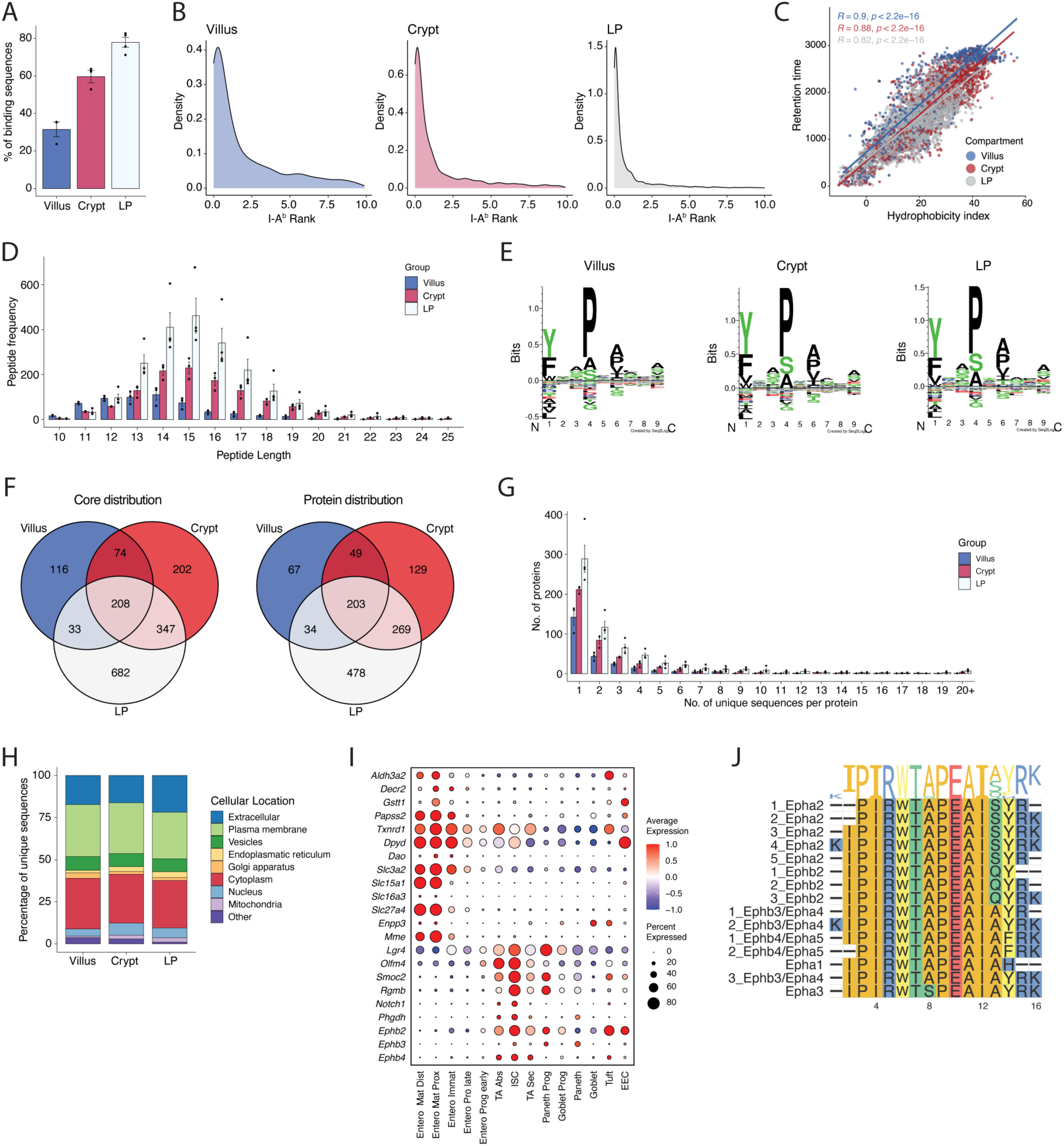
Quality control of the immunopeptidomics data. Analysis of MHC II-eluted peptides from three SI compartments: villus and crypt (*n* = 3 biological replicates), and LP (*n* = 4 biological replicates). **(A)** Percentage of peptides predicted to bind the I-A^b^ allele by compartment. Predictions were calculated by NetMHCIIpan v4.3, rank ≤ 10% binding cutoff ^36^ (**Methods**). Bars show mean ± SEM. **(B)** Density plots of NetMHCIIpan ranks for predicted I-A^b^ binding peptides (rank ≤ 10%, post-filtering), by compartment. **(C)** Scatter plot of peptide retention time (RT) versus hydrophobicity index for villus, crypt, and LP immunopeptides (**Methods**). **(D)** Length distribution of identified peptides by compartment (mean ± SEM). **(E)** Sequence motifs of peptide cores from villus (left), crypt (mid), and LP (right) compartments. Peptide sequences were clustered with the GibbsCluster 2.0 server ^62^ and generated with Seq2Logo 2.0 ^63^. **(F)** Venn diagrams of peptide-derived cores (left) and peptide-matched proteins (right) across villus, crypt, and LP compartments. Counts are pooled across biological replicates after filtering. (**G)** Distribution of unique peptides per source protein by compartment. **(H)** Subcellular localization of peptide source proteins by compartment. Bars show the percentage of unique MHC II-bound peptides mapped to proteins and annotated to the subcellular locations (UniProt). Percentages were calculated per compartment from unique peptides; proteins are assigned to their primary UniProt “Subcellular location” term (remaining terms grouped as “Other”). **(I)** Comparison of immunopeptide-specific presentation and RNA-specific expression. Dot plot of gene expression derived from the scRNA-seq data for proteins matched to compartment-enriched peptide cores. Columns represent annotated IEC subsets; rows show selected genes. Dot size indicates the fraction of cells in each cluster with detectable expression; dot color indicates mean expression (z-score). **(J)** Sequence alignment of class II peptides belonging to different Ephrin receptors by ggmsa v1.0.2 R package ^64^. Identified sequences were pooled across intestinal compartments.

**Figure S5.**
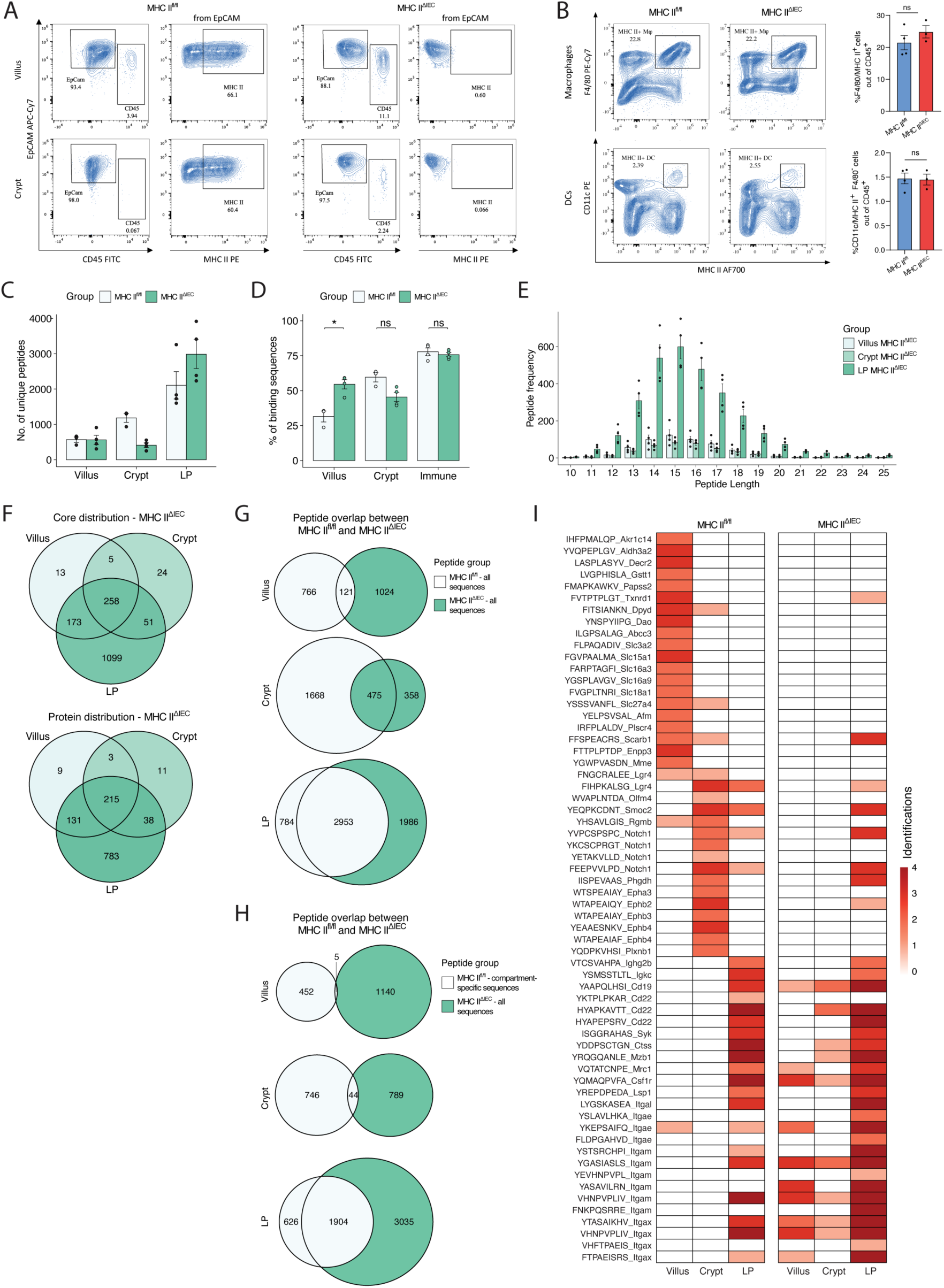
Exploring the MHC II-peptidome under epithelial MHC II knockout. **(A-B)** Flow cytometry (FC) analysis of intestinal compartments following epithelial MHC II deletion (MHC II^ΔIEC^). **(A)** Representative quality control FC plots show MHC II deletion in EpCAM⁺ CD45⁻ villus and crypt IECs of MHC II^ΔIEC^ compared to control MHC II^fl/fl^ IECs. **(B)** Percentage of MHC II⁺ macrophages (upper panel) and MHC II⁺ dendritic cells (lower panel) among CD45⁺ LP immune cells in MHC II^fl/fl^ and MHC II^ΔIEC^ mice. Representative flow cytometry plots are shown (left panel). Quantification of MHC II^+^ cells from MHC II^fl/fl^ and MHC II^ΔIEC^ mice (right panel). Data are shown as mean ± SEM; statistical analysis was performed using two-tailed unpaired *t-test* with Welch’s correction (*n* = 4 mice of MHC II^fl/fl^, *n* = 3 mice of MHC II^ΔIEC^). **(C)** Number of unique peptide sequences per compartment in MHC II^fl/fl^ vs MHC II^ΔIEC^ mice, after filtering. **(D)** Percentage of peptides predicted to bind the I-Ab allele in MHC II^fl/fl^ compared to MHC II^ΔIEC^ mice by compartment according to NetMHCIIpan v4.3 with rank ≤ 10% as binder cutoff. Shown as mean ± SEM; statistical analysis was performed by multiple unpaired t-tests with Holm-Šídák correction (α = 0.05). \**P* < 0.05. **(E)** Length distribution of identified peptides in MHC II^ΔIEC^ by compartment. Bar plot with mean ± SEM indicates the number of unique peptides per sample. **(F)** Overlap of peptide-derived cores (upper panel) and peptide-matched source proteins (lower panel) in the villus, crypt, and LP immune cells compartment of MHC II^ΔIEC^ mice. Counts are pooled across replicates. **(G-H)** Peptide overlap between MHC II^fl/fl^ and MHC II^ΔIEC^ mice in the villus, crypt, and LP compartments. Counts are pooled across replicates. **(G)** Comparison of all identified sequences between MHC II^fl/fl^ and MHC II^ΔIEC^ mice. **(H)** Overlap between compartment-specific sequences of MHC II^fl/fl^ mice and all identified sequences of MHC II^ΔIEC^ mice. (**I**) Epithelial compartment-depleted peptide cores. Each tile indicates the number of biological replicates in which the peptide core was detected in the given compartment and condition.

**Figure S6.**
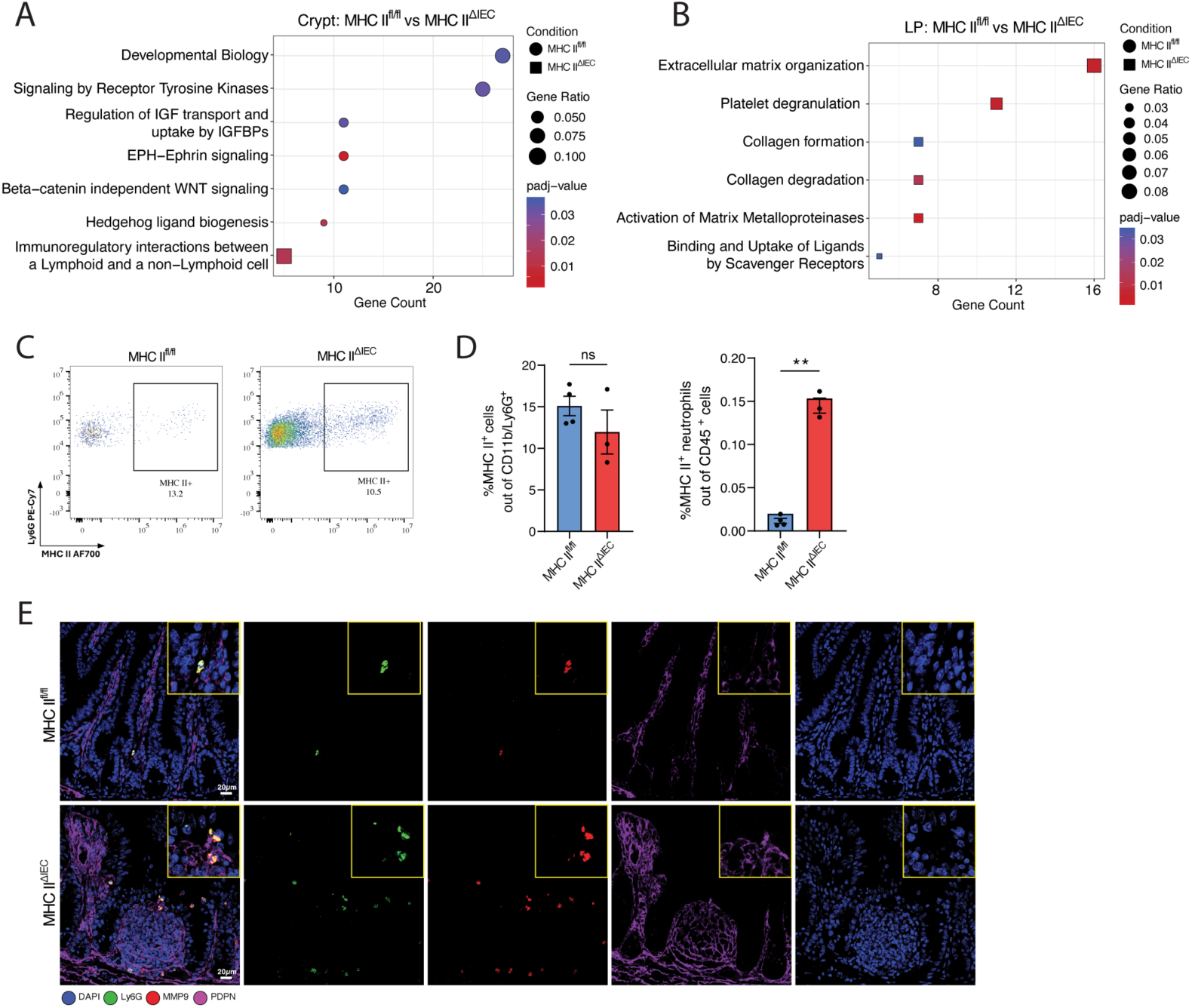
ECM remodeling and immune infiltration under MHC II^ΔIEC^. **(A-B)** Reactome pathway enrichment analysis of unique source proteins from the crypt **(A)** and LP **(B)** immunopeptidome in MHC II^ΔIEC^ compared to MHC II^fl/fl^ mice by the ReactomePA v1.16.2 R package ^58^. Significance was assessed using Benjamini-Hochberg correction, *q* < 0.05. **(C-D)** Flow cytometry analysis of neutrophil changes in MHC II^ΔIEC^ mice. **(C)** Representative FC plots showing an increase of Ly6G^+^ CD11b^+^ CD45^+^ neutrophils in MHC II^ΔIEC^ mice. **(D)** Percentage of MHC II⁺ cells out of neutrophils (left) and percentage of MHC II⁺ neutrophils out of CD45^+^ cells (right) in MHC II^fl/fl^ compared to MHC II^ΔIEC^ mice. Statistical analysis was performed using a two-tailed unpaired t-test with Welch’s correction; *n* = 4 mice MHC II^fl/fl^, *n* = 3 mice MHC II^ΔIEC^. \*\**P* < 0.01. (**E**) Colocalization of Lyz1 (green) and MMP9 (red) markers of neutrophils in the distal SI. Fibroblasts were stained with PDPN (purple). Nuclei stained with DAPI (blue). Scale bar: 20µm.

**Figure S7.**
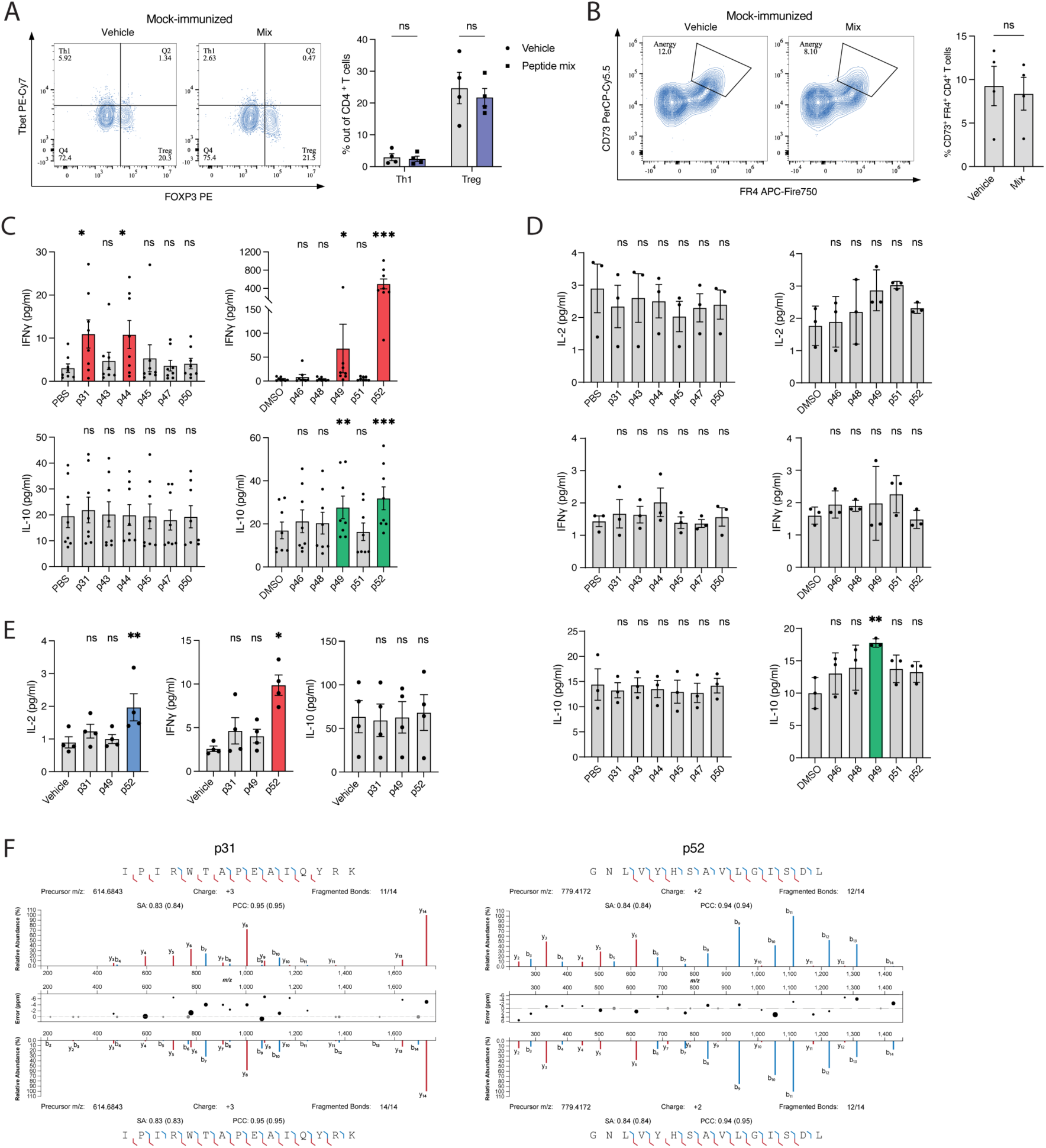
Characterizing the immune properties of ISC class II-peptides. **(A,B)** CD4^+^ T cells expansion upon antigen restimulation. Splenocytes of WT mice immunized only with adjuvant were used. Splenocytes were cultured with a peptide mix (2 μg/ml each) for 24h. **(A)** Frequency of Th1 (T-bet^+^ FOXP3^-^) and Treg (FOXP3^+^) cells out of activated CD4^+^ T cells (CD45^+^ CD3^+^ CD4^+^ CD44^+^) was analyzed. Representative flow cytometry plots were shown. Statistical analysis by multiple paired t-tests with Holm-Šídák correction (α = 0.05), mean ± SEM (*n* = 4 mice per group). **(B)** Frequency of CD73^+^/FR4^+^ T cells out of activated non-Treg CD4^+^ T cells (CD45^+^ CD3^+^ CD4^+^ CD44^+^ FOXP3^-^). Representative flow cytometry plots were shown. Statistical analysis by the two-tailed paired t-tests, mean ± SEM (*n* = 4 mice). **(C,D)** Splenocyte response was profiled after immunization with a cocktail of 11 ISC peptides + adjuvant **(C)** or with adjuvant alone **(D)**. Peptides were split into two groups: PBS-soluble and DMSO-soluble; each group had its volume-matched vehicle control (PBS or DMSO, respectively). IL-2, IFNψ, and IL-10 secretion were measured in response to individual ISC-specific class II peptides (20 μg/ml) by CBA assay. Matched (repeated-measures) data were analyzed with a Friedman test, followed by Dunn’s multiple-comparisons test comparing each peptide to the common control (two-sided; PBS- and DMSO-groups were analyzed separately). Reported *P* values are multiplicity-adjusted across all peptide-vs-control tests. Data are mean ± SEM, *n* = 8 mice **(C)** and *n* =3 mice **(D)**. \**P* < 0.05, \*\**P* < 0.01, \*\*\**P* < 0.001, ns = non-significant. **(E)** Immune cells from LP were profiled after immunization with a cocktail of 11 ISC peptides + adjuvant. IL-2, IFNψ, and IL-10 secretion were measured in response to individual ISC-specific class II peptides (20 μg/ml) by CBA assay. Responses to selected peptides were compared with volume-matched vehicle controls. Matched (repeated-measures) data were analyzed with a Friedman test, followed by Dunn’s multiple-comparisons test comparing each peptide to the common control (two-sided). Reported P values are multiplicity-adjusted across all peptide-vs-control tests. Data are mean ± SEM, n = 4 mice. \**P* < 0.05, \*\**P* < 0.01. **(F)** Mirror plots depict the peptide spectra matches of p31 (left) and p52 (right) peptides between experimentally identified (top) and synthetic (bottom) peptides (PCC: Pearson correlation coefficient; SA: Spectrum angle).

**Figure S8.**
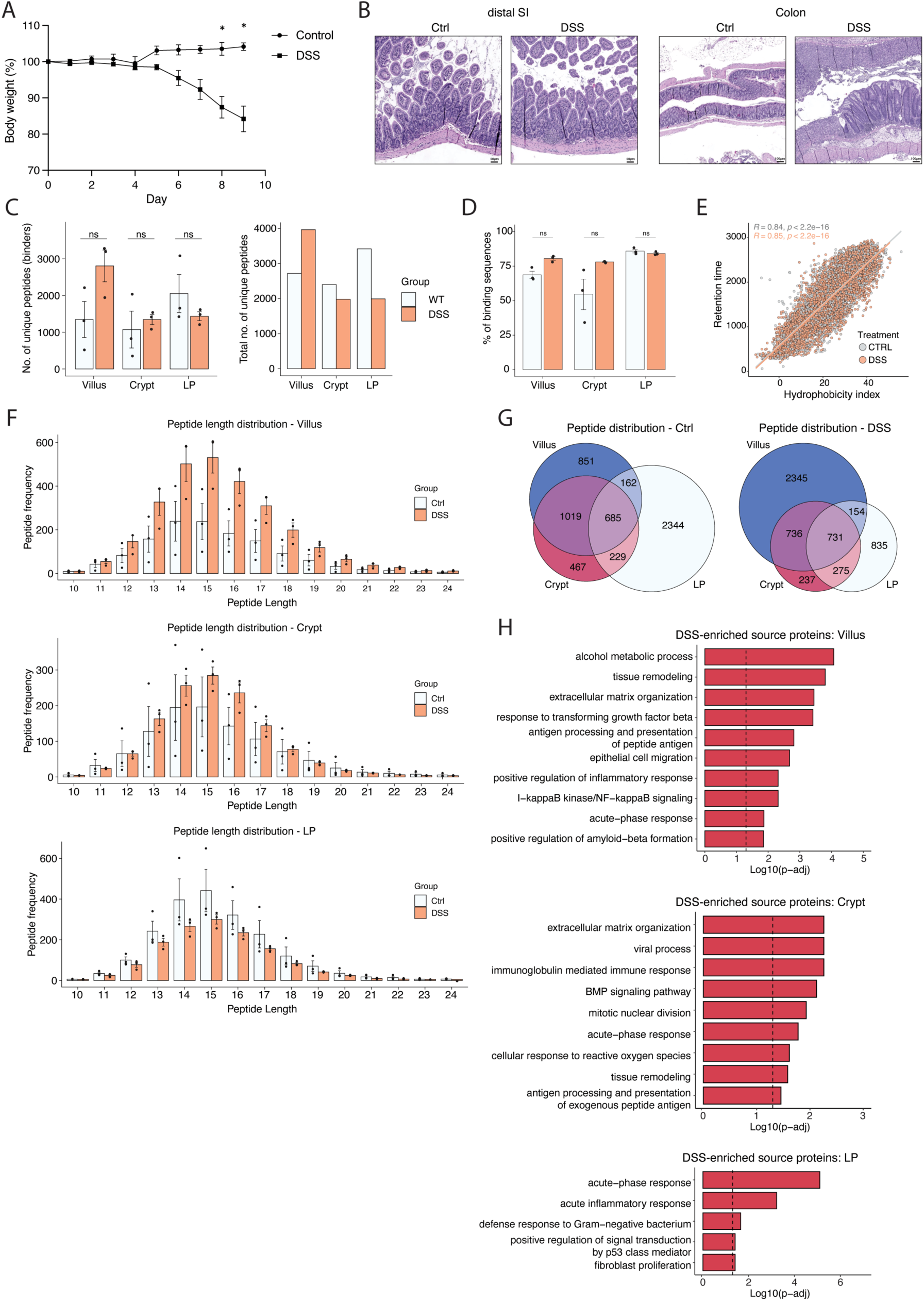
Identification of gut injury class II immunopeptides. (**A**) Body weight change over the course of DSS treatment. Shown as mean ± SEM (Control *n* = 3 mice, DSS *n* = 5 mice); statistical analysis by mixed-effects model (REML) with time and treatment as fixed effects and mouse as a random effect, followed by Sidak-corrected post hoc comparisons between groups at each time point. \**P* < 0.05. (**B**) Representative images of H&E staining of distal SI sections (left panel, scale bar 50 µm) and colon (right panel, scale bar 100 µm) from control (Ctrl) vs DSS-treated mice. (**C**-**H**) Immunopeptidomics analysis of the villus, crypt, and LP compartments from control vs DSS-treated mice (*n* = 3 mice per group). (**C**) Number of unique MHC II-bound peptides per compartment. Left panel, mean ± SEM with per-sample points overlaid; right panel, total number of unique sequences per compartment. (**D**) Percentage of peptides predicted to bind the I-A^b^ allele by compartment and condition. Prediction done by NetMHCIIpan v4.3 ^36^ using rank ≤ 10% as binder cut-off. Statistical analysis by multiple unpaired t-tests with Holm-Šídák correction (α = 0.05), mean ± SEM. (**E**) Scatter plot of retention time (RT) vs hydrophobicity index for control- vs DSS-identified immunopeptides. Immunopeptides were combined across all compartments after filtering. (**F**) Length distribution of identified unique peptides after filtering, in each analyzed compartment (top, mid, and bottom); mean ± SEM. (**G**) Distribution of the identified peptides across intestinal compartments of control vs DSS-treated mice; counts pooled across replicates. (**H**) GO analysis for proteins enriched in DSS compared to control per compartment (villus, crypt, LP). Analysis done by clusterProfiler R package with Benjamini-Hochberg FDR (**Methods**); bars are log₁₀(*q* value) with the dashed line at *q* = 0.05.

## References

1. Pishesha, N., Harmand, T.J., and Ploegh, H.L. (2022). A guide to antigen processing and presentation. Nat. Rev. Immunol. 22, 751–764.

2. Rock, K.L., Reits, E., and Neefjes, J. (2016). Present yourself! By MHC class I and MHC class II molecules. Trends Immunol. 37, 724–737.

3. Kedmi, R., and Littman, D.R. (2024). Antigen-presenting cells as specialized drivers of intestinal T cell functions. Immunity 57, 2269–2279.

4. Cummings, R.J., Barbet, G., Bongers, G., Hartmann, B.M., Gettler, K., Muniz, L., Furtado, G.C., Cho, J., Lira, S.A., and Blander, J.M. (2016). Different tissue phagocytes sample apoptotic cells to direct distinct homeostasis programs. Nature 539, 565–569.

5. Coombes, J.L., Siddiqui, K.R.R., Arancibia-Cárcamo, C.V., Hall, J., Sun, C.-M., Belkaid, Y., and Powrie, F. (2007). A functionally specialized population of mucosal CD103+ DCs induces Foxp3+ regulatory T cells via a TGF-β– and retinoic acid–dependent mechanism. J. Exp. Med. 204, 1757–1764.

6. Mazzini, E., Massimiliano, L., Penna, G., and Rescigno, M. (2014). Oral Tolerance Can Be Established via Gap Junction Transfer of Fed Antigens from CX3CR1+ Macrophages to CD103+ Dendritic Cells. Immunity 40, 248–261.

7. Fu, L., Upadhyay, R., Pokrovskii, M., Chen, F.M., Romero-Meza, G., Griesemer, A., and Littman, D.R. (2025). PRDM16-dependent antigen-presenting cells induce tolerance to gut antigens. Nature 642, 756–765.

8. Rodrigues, P.F., Wu, S., Trsan, T., Panda, S.K., Fachi, J.L., Liu, Y., Du, S., de Oliveira, S., Antonova, A.U., Khantakova, D., et al. (2025). Rorγt-positive dendritic cells are required for the induction of peripheral regulatory T cells in response to oral antigens. Cell 188, 2720–2737.e22.

9. Zhou, Y.D., Komnick, M.R., and Esterházy, D. (2026). Dendritic cells in the gastrointestinal system: Division of labor, plasticity, and niche-specific adaptation. Immunol. Rev. 337, e70090.

10. Heuberger, C., Pott, J., and Maloy, K.J. (2021). Why do intestinal epithelial cells express MHC class II? Immunology 162, 357–367.

11. Wosen, J.E., Mukhopadhyay, D., Macaubas, C., and Mellins, E.D. (2018). Epithelial MHC class II expression and its role in antigen presentation in the gastrointestinal and respiratory tracts. Front. Immunol. 9, 2144.

12. Malik, A., Sharma, D., Aguirre-Gamboa, R., McGrath, S., Zabala, S., Weber, C., and Jabri, B. (2023). Epithelial IFNγ signalling and compartmentalized antigen presentation orchestrate gut immunity. Nature 623, 1044–1052.

13. He, K., Wan, T., Wang, D., Hu, J., Zhou, T., Tao, W., Wei, Z., Lu, Q., Zhou, R., Tian, Z., et al. (2023). Gasdermin D licenses MHCII induction to maintain food tolerance in small intestine. Cell 186, 3033–3048.e20.

14. Biton, M., Haber, A.L., Rogel, N., Burgin, G., Beyaz, S., Schnell, A., Ashenberg, O., Su, C.-W., Smillie, C., Shekhar, K., et al. (2018). T helper cell cytokines modulate intestinal stem cell renewal and differentiation. Cell 175, 1307–1320.e22.

15. Tuganbaev, T., Mor, U., Bashiardes, S., Liwinski, T., Nobs, S.P., Leshem, A., Dori-Bachash, M., Thaiss, C.A., Pinker, E.Y., Ratiner, K., et al. (2020). Diet diurnally regulates small intestinal microbiome-epithelial-immune homeostasis and enteritis. Cell 182, 1441–1459.e21.

16. Eshleman, E.M., Shao, T.-Y., Woo, V., Rice, T., Engleman, L., Didriksen, B.J., Whitt, J., Haslam, D.B., Way, S.S., and Alenghat, T. (2023). Intestinal epithelial HDAC3 and MHC class II coordinate microbiota-specific immunity. J. Clin. Invest. 133. 10.1172/JCI162190.

17. Koyama, M., Mukhopadhyay, P., Schuster, I.S., Henden, A.S., Hülsdünker, J., Varelias, A., Vetizou, M., Kuns, R.D., Robb, R.J., Zhang, P., et al. (2019). MHC class II antigen presentation by the intestinal epithelium initiates graft-versus-host disease and is influenced by the Microbiota. Immunity 51, 885–898.e7.

18. Jamwal, D.R., Laubitz, D., Harrison, C.A., Figliuolo da Paz, V., Cox, C.M., Wong, R., Midura-Kiela, M., Gurney, M.A., Besselsen, D.G., Setty, P., et al. (2020). Intestinal epithelial expression of MHCII determines severity of chemical, T-cell-induced, and infectious colitis in mice. Gastroenterology 159, 1342–1356.e6.

19. Westendorf, A.M., Bruder, D., Hansen, W., and Buer, J. (2006). Intestinal epithelial antigen induces CD4+ T cells with regulatory phenotype in a transgenic autoimmune mouse model. Ann. N. Y. Acad. Sci. 1072, 401–406.

20. Westendorf, A.M., Fleissner, D., Groebe, L., Jung, S., Gruber, A.D., Hansen, W., and Buer, J. (2009). CD4+Foxp3+ regulatory T cell expansion induced by antigen-driven interaction with intestinal epithelial cells independent of local dendritic cells. Gut 58, 211–219.

21. Unanue, E.R., Turk, V., and Neefjes, J. (2016). Variations in MHC class II antigen processing and presentation in health and disease. Annu. Rev. Immunol. 34, 265–297.

22. Cho, J.H., and Feldman, M. (2015). Heterogeneity of autoimmune diseases: pathophysiologic insights from genetics and implications for new therapies. Nat. Med. 21, 730–738.

23. Kim, M.W., Gao, W., Lichti, C.F., Gu, X., Dykstra, T., Cao, J., Smirnov, I., Boskovic, P., Kleverov, D., Salvador, A.F.M., et al. (2025). Endogenous self-peptides guard immune privilege of the central nervous system. Nature 637, 176–183.

24. Krienke, C., Kolb, L., Diken, E., Streuber, M., Kirchhoff, S., Bukur, T., Akilli-Öztürk, Ö., Kranz, L.M., Berger, H., Petschenka, J., et al. (2021). A noninflammatory mRNA vaccine for treatment of experimental autoimmune encephalomyelitis. Science 371, 145–153.

25. Abelin, J.G., Keskin, D.B., Sarkizova, S., Hartigan, C.R., Zhang, W., Sidney, J., Stevens, J., Lane, W., Zhang, G.L., Eisenhaure, T.M., et al. (2017). Mass spectrometry profiling of HLA-associated peptidomes in mono-allelic cells enables more accurate Epitope prediction. Immunity 46, 315–326.

26. Abelin, J.G., Harjanto, D., Malloy, M., Suri, P., Colson, T., Goulding, S.P., Creech, A.L., Serrano, L.R., Nasir, G., Nasrullah, Y., et al. (2019). Defining HLA-II ligand processing and binding rules with mass spectrometry enhances cancer Epitope prediction. Immunity 51, 766–779.e17.

27. Purcell, A.W., Ramarathinam, S.H., and Ternette, N. (2019). Mass spectrometry-based identification of MHC-bound peptides for immunopeptidomics. Nat. Protoc. 14, 1687–1707.

28. Arshad, S., Cameron, B., and Joglekar, A.V. (2025). Immunopeptidomics for autoimmunity: unlocking the chamber of immune secrets. NPJ Syst. Biol. Appl. 11, 10.

29. Wan, X., Vomund, A.N., Peterson, O.J., Chervonsky, A.V., Lichti, C.F., and Unanue, E.R. (2020). The MHC-II peptidome of pancreatic islets identifies key features of autoimmune peptides. Nat. Immunol. 21, 455–463.

30. Thelemann, C., Eren, R.O., Coutaz, M., Brasseit, J., Bouzourene, H., Rosa, M., Duval, A., Lavanchy, C., Mack, V., Mueller, C., et al. (2014). Interferon-γ induces expression of MHC class II on intestinal epithelial cells and protects mice from colitis. PLoS One 9, e86844.

31. Barker, N., van Es, J.H., Kuipers, J., Kujala, P., van den Born, M., Cozijnsen, M., Haegebarth, A., Korving, J., Begthel, H., Peters, P.J., et al. (2007). Identification of stem cells in small intestine and colon by marker gene Lgr5. Nature 449, 1003–1007.

32. Haber, A.L., Biton, M., Rogel, N., Herbst, R.H., Shekhar, K., Smillie, C., Burgin, G., Delorey, T.M., Howitt, M.R., Katz, Y., et al. (2017). A single-cell survey of the small intestinal epithelium. Nature 551, 333–339.

33. Zheng, D., Liwinski, T., and Elinav, E. (2020). Interaction between microbiota and immunity in health and disease. Cell Res. 30, 492–506.

34. Brown, C.C., and Rudensky, A.Y. (2023). Spatiotemporal regulation of peripheral T cell tolerance. Science 380, 472–478.

35. Kalaora, S., and Samuels, Y. (2019). Cancer exome-based identification of tumor Neo-antigens using mass spectrometry. Methods Mol. Biol. 1884, 203–214.

36. Nilsson, J.B., Kaabinejadian, S., Yari, H., Kester, M.G.D., van Balen, P., Hildebrand, W.H., and Nielsen, M. (2023). Accurate prediction of HLA class II antigen presentation across all loci using tailored data acquisition and refined machine learning. Sci. Adv. 9, eadj6367.

37. Bassani-Sternberg, M., Bräunlein, E., Klar, R., Engleitner, T., Sinitcyn, P., Audehm, S., Straub, M., Weber, J., Slotta-Huspenina, J., Specht, K., et al. (2016). Direct identification of clinically relevant neoepitopes presented on native human melanoma tissue by mass spectrometry. Nat. Commun. 7, 13404.

38. Fugmann, T., Sofron, A., Ritz, D., Bootz, F., and Neri, D. (2017). The MHC class II immunopeptidome of lymph nodes in health and in chemically induced colitis. J. Immunol. 198, 1357–1364.

39. Dengjel, J., Schoor, O., Fischer, R., Reich, M., Kraus, M., Müller, M., Kreymborg, K., Altenberend, F., Brandenburg, J., Kalbacher, H., et al. (2005). Autophagy promotes MHC class II presentation of peptides from intracellular source proteins. Proc. Natl. Acad. Sci. U. S. A. 102, 7922–7927.

40. Adamopoulou, E., Tenzer, S., Hillen, N., Klug, P., Rota, I.A., Tietz, S., Gebhardt, M., Stevanovic, S., Schild, H., Tolosa, E., et al. (2013). Exploring the MHC-peptide matrix of central tolerance in the human thymus. Nat. Commun. 4, 2039.

41. Holmberg, J., Genander, M., Halford, M.M., Annerén, C., Sondell, M., Chumley, M.J., Silvany, R.E., Henkemeyer, M., and Frisén, J. (2006). EphB receptors coordinate migration and proliferation in the intestinal stem cell niche. Cell 125, 1151–1163.

42. Migulina, N., de Hilster, R.H.J., Bartel, S., Vedder, R.H.J., van den Berge, M., Nagelkerke, A., Timens, W., Harmsen, M.C., Hylkema, M.N., Brandsma, C.-A., et al. (2024). 3-D culture of human lung fibroblasts decreases proliferative and increases extracellular matrix remodeling genes. Am. J. Physiol. Cell Physiol. 326, C177–C193.

43. Lin, M., Jackson, P., Tester, A.M., Diaconu, E., Overall, C.M., Blalock, J.E., and Pearlman, E. (2008). Matrix metalloproteinase-8 facilitates neutrophil migration through the corneal stromal matrix by collagen degradation and production of the chemotactic peptide Pro-Gly-Pro. Am. J. Pathol. 173, 144–153.

44. Opdenakker, G., Vermeire, S., and Abu El-Asrar, A. (2022). How to place the duality of specific MMP-9 inhibition for treatment of inflammatory bowel diseases into clinical opportunities? Front. Immunol. 13. 10.3389/FIMMU.2022.983964.

45. Adir, I., Sochen, C., Menachem, A.H., Lebon, S., Toval, B., Holiar, V., Davidzohn, N., Salame, T.-M., Rosenhek-Goldian, I., Savickas, S., et al. (2025). Persistent ECM Scarring Reprograms Intestinal Stem Cells to Drive Chronic Inflammation. Immunology.

46. Muñoz, J., Stange, D.E., Schepers, A.G., van de Wetering, M., Koo, B.-K., Itzkovitz, S., Volckmann, R., Kung, K.S., Koster, J., Radulescu, S., et al. (2012). The Lgr5 intestinal stem cell signature: robust expression of proposed quiescent “+4” cell markers. EMBO J. 31, 3079–3091.

47. Shibaki, A., and Katz, S.I. (2002). Induction of skewed Th1/Th2 T-cell differentiation via subcutaneous immunization with Freund’s adjuvant: Cellular immune responses via CFA immunization. Exp. Dermatol. 11, 126–134.

48. Pérez-Cruz, M., Iliopoulou, B.P., Hsu, K., Wu, H.-H., Erkers, T., Swaminathan, K., Tang, S.-W., Bader, C.S., Kambham, N., Xie, B., et al. (2022). Immunoregulatory effects of RGMb in gut inflammation. Front. Immunol. 13, 960329.

49. Shi, Y., Zhong, L., Li, Y., Chen, Y., Feng, S., Wang, M., Xia, Y., and Tang, S. (2021). Repulsive guidance molecule b deficiency induces gut Microbiota dysbiosis and increases the susceptibility to intestinal inflammation in mice. Front. Microbiol. 12, 648915.

50. Chen, R., Chen, Q., Zheng, J., Zeng, Z., Chen, M., Li, L., and Zhang, S. (2023). Serum amyloid protein A in inflammatory bowel disease: from bench to bedside. Cell Death Discov. 9, 154.

51. Peterson, L.W., and Artis, D. (2014). Intestinal epithelial cells: regulators of barrier function and immune homeostasis. Nat. Rev. Immunol. 14, 141–153.

52. Iberg, C.A., Jones, A., and Hawiger, D. (2017). Dendritic cells as inducers of peripheral tolerance. Trends Immunol. 38, 793–804.

53. Coombes, J.L., and Powrie, F. (2008). Dendritic cells in intestinal immune regulation. Nat. Rev. Immunol. 8, 435–446.

54. Scott, C.L., Aumeunier, A.M., and Mowat, A.M. (2011). Intestinal CD103+ dendritic cells: master regulators of tolerance? Trends Immunol. 32, 412–419.

55. Zhou, T., Zhang, G., Wu, C., Wan, T., and Zhu, S. (2025). Type 1 Regulatory T cells induced by intestinal epithelial cells respond to food antigen. bioRxiv. 10.1101/2025.07.23.666039.

56. Lee, J.-Y., Hall, J.A., Kroehling, L., Wu, L., Najar, T., Nguyen, H.H., Lin, W.-Y., Yeung, S.T., Silva, H.M., Li, D., et al. (2020). Serum amyloid A proteins induce pathogenic Th17 cells and promote inflammatory disease. Cell 180, 79–91.e16.

57. Kang, K., Huang, C., Li, Y., Umbach, D.M., and Li, L. (2021). CDSeqR: fast complete deconvolution for gene expression data from bulk tissues. BMC Bioinformatics 22, 262.

58. Yu, G., and He, Q.-Y. (2016). ReactomePA: an R/Bioconductor package for reactome pathway analysis and visualization. Mol. Biosyst. 12, 477–479.

59. Love, M.I., Huber, W., and Anders, S. (2014). Moderated estimation of fold change and dispersion for RNA-seq data with DESeq2. Genome Biol. 15, 1–21.

60. Xu, S., Hu, E., Cai, Y., Xie, Z., Luo, X., Zhan, L., Tang, W., Wang, Q., Liu, B., Wang, R., et al. (2024). Using clusterProfiler to characterize multiomics data. Nat. Protoc. 19, 3292–3320.

61. Andreatta, M., and Carmona, S.J. (2021). UCell: Robust and scalable single-cell gene signature scoring. Comput. Struct. Biotechnol. J. 19, 3796.

62. Andreatta, M., Lund, O., and Nielsen, M. (2013). Simultaneous alignment and clustering of peptide data using a Gibbs sampling approach. Bioinformatics 29, 8–14.

63. Thomsen, M.C.F., and Nielsen, M. (2012). Seq2Logo: a method for construction and visualization of amino acid binding motifs and sequence profiles including sequence weighting, pseudo counts and two-sided representation of amino acid enrichment and depletion. Nucleic Acids Res. 40, W281–7.

64. Zhou, L., Feng, T., Xu, S., Gao, F., Lam, T.T., Wang, Q., Wu, T., Huang, H., Zhan, L., Li, L., et al. (2022). Ggmsa: A visual exploration tool for multiple sequence alignment and associated data. Brief. Bioinform. 23. 10.1093/bib/bbac222.

65. Foroutan, M., Bhuva, D.D., Lyu, R., Horan, K., Cursons, J., and Davis, M.J. (2018). Single sample scoring of molecular phenotypes. BMC Bioinformatics 19, 404.

66. Kong, A.T., Leprevost, F.V., Avtonomov, D.M., Mellacheruvu, D., and Nesvizhskii, A.I. (2017). MSFragger: ultrafast and comprehensive peptide identification in mass spectrometry-based proteomics. Nat. Methods 14, 513–520.

67. Weller, C., Bartok, O., McGinnis, C.S., Palashati, H., Chang, T.-G., Malko, D., Shmueli, M.D., Nagao, A., Hayoun, D., Murayama, A., et al. (2025). Translation dysregulation in cancer as a source for targetable antigens. Cancer Cell 43, 823–840.e18.

68. Krokhin, O.V., Craig, R., Spicer, V., Ens, W., Standing, K.G., Beavis, R.C., and Wilkins, J.A. (2004). An improved model for prediction of retention times of tryptic peptides in ion pair reversed-phase HPLC. Mol. Cell. Proteomics 3, 908–919.

69. Virtanen, P., Gommers, R., Oliphant, T.E., Haberland, M., Reddy, T., Cournapeau, D., Burovski, E., Peterson, P., Weckesser, W., Bright, J., et al. (2020). SciPy 1.0: fundamental algorithms for scientific computing in Python. Nat. Methods 17, 261–272.

70. Gessulat, S., Schmidt, T., Zolg, D.P., Samaras, P., Schnatbaum, K., Zerweck, J., Knaute, T., Rechenberger, J., Delanghe, B., Huhmer, A., et al. (2019). Prosit: proteome-wide prediction of peptide tandem mass spectra by deep learning. Nat. Methods 16, 509–518.

